# Universal Dynamic Fitting of Magnetic Resonance Spectroscopy

**DOI:** 10.1101/2023.06.15.544935

**Authors:** William T Clarke, Clémence Ligneul, Michiel Cottaar, I Betina Ip, Saad Jbabdi

## Abstract

Dynamic (2D) magnetic resonance spectroscopy is a collection of techniques where acquisitions of spectra are repeated under varying experimental or physiological conditions. Dynamic MRS comprises a rich set of contrasts, including diffusion-weighted, relaxation-weighted, functional, edited, or hyperpolarized spectroscopy, leading to quantitative insights into multiple physiological or microstructural processes. Conventional approaches to dynamic MRS analysis ignore the shared information between spectra, and instead proceed by independently fitting noisy individual spectra before modelling temporal changes in the parameters. Here we propose a universal dynamic MRS toolbox which allows simultaneous fitting of dynamic spectra of arbitrary type. A simple interface allows information to be shared and precisely modelled across spectra to make inferences on both spectral and dynamic processes. We demonstrate and thoroughly evaluate our approach in three types of dynamic MRS techniques. Simulations of functional and edited MRS are used to demonstrate the advantages of dynamic fitting. Analysis of synthetic functional 1H-MRS data shows a marked decrease in parameter uncertainty as predicted by prior work. Analysis with our tool replicates the results of two previously published studies using the original in vivo functional and diffusion-weighted data. Finally, joint spectral fitting with diffusion orientation models is demonstrated in synthetic data. The toolbox is shared as a fully open-source software with comprehensive documentation, example data, and tutorials.

## Introduction

During dynamic, or 2D, magnetic resonance spectroscopy (MRS), multiple spectra are acquired whilst experimental conditions change. Dynamic changes can be induced deliberately, to sensitise acquisitions to different signal mechanisms. Conditions can also change due to uncontrollable physiological processes, such as structured noise from cardiorespiratory motion or voluntary movement^1^, or due to hardware drift.^2^ In all types of dynamic MRS, the classical processing pipelines start by fitting a spectral model to each transient,^3,4^ or to averages of repeated measurements.^5,6^ They then extract parameters of interest from these fits, usually metabolite concentrations, and analyse or model their changes across experimental conditions. However, MRS is an inherently low signal-to-noise technique compared to proton-MRI, as metabolites occur with concentrations thousands of times lower than water. This means repeated measurements are required, at the detriment of more interesting and informative changes induced experimentally. Simultaneous fitting of all spectra, i.e., dynamic fitting, can mitigate this trade-off, by modelling the effect of changing the experimental conditions and by explicitly sharing relevant information across dynamic spectra.

For example, in spectral editing^7^, two or more spectra are acquired with appropriate changes in the pulse sequence aimed at suppressing the signal around targeted spectral peaks. While these spectra may be affected by different factors that require separate modelling, such as phase shifts,^8^ they share the fact that the underlying metabolite concentrations are unaffected by the pulse sequence. A straightforward dynamic fit can estimate the shared concentrations while independently fitting nuisance factors. A similar logic applies for non-edited spectroscopy, where noisy transients are affected by separate artefacts whilst metabolite concentrations remain constant.^2^ In diffusion-weighted MRS, the apparent concentrations are reduced as a function of the diffusion encoding due to the random motion of metabolites.^9^ Models linking metabolite diffusion to the underlying tissue microstructure can be used to link across dynamic spectra,^10^ thereby imposing a precise structure to help the fitting, particularly when strong diffusion-encoding drastically decreases signal-to-noise. A similar approach can be used in functional MRS, where the experimental manipulation is usually an exogenous stimulus,^11^ which effect on the concentrations and potentially on other parameters such as the linewidth can be explicitly modelled.^12^

In this work we introduce an extension to FSL-MRS^14^ that allows direct fitting of an arbitrary dynamic signal model to multiple spectra simultaneously. FSL-MRS is an end-to-end spectroscopy analysis toolbox embedded in the widely-used FSL neuroimaging analysis environment.^15^ Incorporation of a dynamic model with spectral fitting of multiple signal transients reduces the number of parameters to be estimated from noisy data, and, as demonstrated by *Tal*^13^ and in this work, reduces fitting uncertainty. These enhancements also establish a framework for model selection, as well as robust statistical testing at the group-level, when data are combined across subjects or sessions. These new tools are embedded into a wider spectroscopy analysis (and even wider neuroimaging package) to enable integrated pre-processing of dynamic MRS data. Whilst some similar tools are beginning to emerge,^16^ the toolbox described here is open source, freely available, and importantly, allows arbitrary model flexibility for arbitrary types of dynamic MRS experiments.

Here, we demonstrate uses across in vivo and simulated data to evidence the suitability of the tool for a three MRS contrasts which represent potential common use cases for dynamic fitting. These contrasts are:

1. spectral editing of the metabolite GABA (MEGA-PRESS^8^), using synthetic data,
2. functional MRS (fMRS) measured during visual stimulation, using synthetic and in vivo data, and,
3. diffusion-weighted MRS (dMRS), using synthetic and in vivo data.

We replicate the results of Tal^13^ showing the value of dynamic fitting in improving parameter estimation and uncertainty in a general framework, before extending the analysis to simulation of real-world fMRS data. We further show improvements in errors with spectral editing, validate the accuracy of the implementation in fMRS, and demonstrate how the toolset can be used to mitigate confounds and unlock new measurement approaches. As a further validation, we replicate the results of two in vivo studies (fMRS and dMRS respectively) using their original data. The proposed toolset is released as part of an open-source software package (FSL-MRS), free for academic use. All code and data used in this work is available openly, online.

## Methods

### Model

We describe the evolution of model parameters as a ‘time dependence’, irrespective of how experimental conditions change. We use the linear combination spectral fitting model of FSL-MRS,^14,17^ modified to allow time dependence for all model parameters:

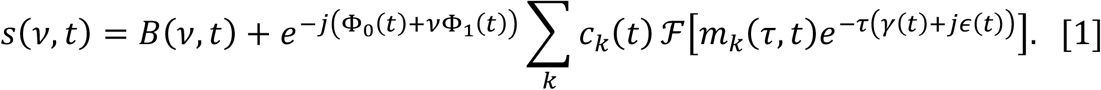

Where the time dependence of the concentration *c*_*k*_, lineshape γ, shift ϵ, phase Φ, and baseline *B* parameters are specified in an editable, Python language, configuration file. Each of the parameters can have their own time dependent behaviour. Parameters may be fixed across all time-points, estimated per timepoint, or constrained to an analytical or numerical model across time. When combined with the ability to specify a linear combination basis set of spectra per time-point, the configuration file approach enables the description of many different types of dynamic MRS. Arbitrary dynamic models may be specified for each of the time-dependent variables in Equation [1]. Each dynamic model can come with its own set of dynamic parameters, which are estimated alongside the spectral parameters using all the data at once. More details on model initialisation and fitting can be found in the Supplementary Material.

### Dynamic Model Specification

The joint spectral-dynamic model is specified through a user-defined configuration file which details the choices of dynamic fitting and the associated dynamic parameters. The user can also specify a time variable input, which contains information about the experimental design leading to dynamic changes. For example, the b-values and gradient directions for dMRS, or a stimulus design matrix for fMRS. The core spectral fitting model is then specified in the same way as a normal linear combination model, as in non-dynamic FSL-MRS fitting.

The Configuration File is simply a Python language text file containing three sections:

1. the time-dependent behaviour of each spectral parameter: fixed, fully variable, or model-constrained,
2. fitting bounds for each free parameter (optional),
3. arbitrary dynamic-model definitions as Python-language functions.

Time-dependence may be defined for a sub-group of parameters, e.g., metabolite concentrations or FSL-MRS “metabolite-groups” (which link frequency shift and line broadening parameters). Dynamic model functions must also provide analytical or numerical gradient definitions in the configuration file. Example configuration files are included in Supporting Figure S1.

### Higher-level / Group Analysis

FSL-MRS implements python scripting (`fsl_mrs.utils.fmrs_tools`) and command line (`fmrs_stats`) interfaces to carry out higher-level or group-level analysis of the results of the dynamic fitting. These tools constitute a Python wrapper around the FSL tool FLAMEO, which implements multilevel linear modelling for group analysis using Bayesian inference.^18^ The tools allow the formation of both first level linear contrasts and high level (group) contrasts, and includes the ability to combine metabolites when the underlying first-level dynamic model is linear.

### Software

Our universal dynamic fitting toolbox is implemented as part of the FSL-MRS spectroscopic analysis package (part of the FMRIB Software Library, FSL^15^), available free of charge for academic use, and published as open-source. Dynamic fitting may be run using interactive or scripted Python coding environments (using the sub-package *fsl_mrs*.*dynamic*) or by using the command-line scripts *fsl_mrs_dynamic*, and *fmrs_stats*. Documentation is provided alongside that of FSL-MRS, in the source-code repository and at *fsl-mrs*.*com*.

FSL-MRS is open-source, with code available online at git.fmrib.ox.ac.uk/fsl/fsl_mrs. Version 2.1.0 of FSL-MRS was used, permanently available at Reference 19. All code and data used in generating this manuscript are available online at git.fmrib.ox.ac.uk/wclarke/fsl-mrs-dynamic-fitting (#c52d6021f3bb64dc89daa80344dcc76c3fa74c2c), and permanently available at References 20 and 21.

### Approach

In this work we use a series of case studies to explore analysis of common dynamic MRS contrasts that benefit from a dynamic fitting approach. There are six case studies, each explores dynamic fitting with a particular contrast, whilst either validating the approach and toolset, or highlighting an advantage over current processing approaches. Each case study is presented with methods and results in the same subsection. The six case studies are:

1. **fMRS: replication and extension of Tal’s monograph**. – shows improved accuracy and precision of correlated parameters in fMRS,
2. **Edited-MRS: improved estimation of [GABA]** – further reveals improved fitting error in metabolite concentrations when dynamic fitting is used,
3. **fMRS: simulated analysis and group statistics** – demonstrates how an analysis of visual stimulation fMRS can be performed, through to group-level statistics,
4. **fMRS: in vivo confound mitigation** – explores the ability to dynamically model blood-oxygen-level-dependent signal as a confound in the fMRS model in real data,
5. **dMRS: multi-direction diffusion encoding** – shows how dynamic fitting allows higher dynamic encoding resolution than would otherwise be restricted by SNR,
6. **dMRS: in vivo validation** – demonstrates a full study analysis using analytical diffusion signal representations, with different models applied to different metabolites.

Our approach is to show that dynamic fitting reduces error when model parameters are correlated, as predicted by Tal (and shown in CS1 & CS2). We also show that FSL-MRS dynamic fitting advances the analysis approach by either: providing a robust statistical framework (CS3), mitigating confounds (CS4 & CS6), or extending the available acquisition approaches (CS5). CS3 can also be used as a fully featured toolset demonstration, and the results as an implementation validation of the tool.

## Case studies

### CS1. Functional MRS: replication, and extension of Tal (Reference 13)

Recently the advantages of dynamic fitting of 2D data (also called spectral-temporal fitting) were demonstrated theoretically and numerically.^13^ In this work we replicated these results using the software framework of FSL-MRS and extended the simulations from toy (two resonance) examples to realistic 1H-fMRS data, containing many overlapping spectral resonances. Functional MRS temporally resolves MRS to detect changes in neurochemical concentrations (or metabolite visibility), induced by external sensory stimulus or otherwise evoked neural activity.^11^ It is analogous to functional MRI (fMRI).

The first simulation implements Reference 13’s toy example. It uses 64 repetitions of a spectrum containing two Lorentzian peaks at defined, but variable separation (Supporting Figure S3). For the central half (32 repetitions) one peak increased in amplitude by 20%, the other peak remained constant throughout (Supporting Figure S2). This toy simulation of fMRS was fitted using the FSL-MRS dynamic approach implementing a general linear model (GLM) to model the temporal dynamics. A design matrix with two regressors (baseline and rectangular-function stimulation period) was used. For comparison, the data was also fitted using FSL-MRS’s independent spectral fitting routine, and the GLM then fitted to the concentration parameters extracted from the independent spectral fits (as in Figure 1). Each simulation was generated and fit 100 times for each of ten peak separations and three different SNR levels. Fitting was carried out as in the original publication of *Tal*, with two unlinked peaks, ‘Free’, and using the normal FSL-MRS fitting approach, with fixed frequency offsets and linked linewidths, ‘Linked’.

**Figure 1.**
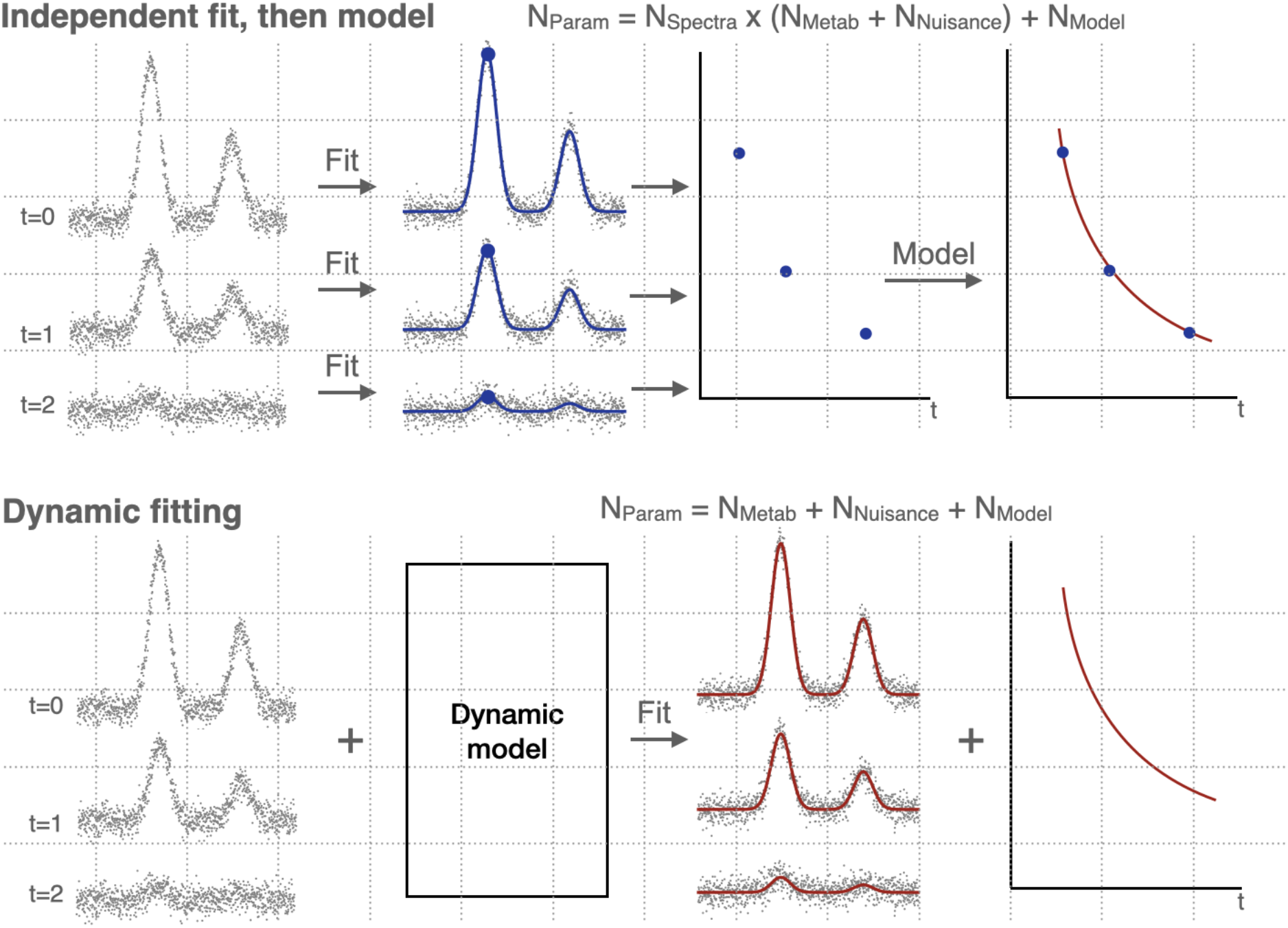
Typical current independent fitting of dynamic data vs. proposed dynamic fitting. The typically used approach in fitting a model to dynamic MRS data (top) is to model the changing parameters after an independent spectral fitting stage (where each spectrum is treated independently). The proposed approach (and as examined by Tal^13^) is to simultaneously fit a spectral and dynamic model. This is known as dynamic, “2D”, or spectral-temporal fitting. This approach reduces the number of parameters to fit by allowing estimation of shared model parameters at once. This shared estimation increases the amount of data used to estimate parameters that are expected to be static (or functionally linked) across transients, mitigating the effect of noise which would otherwise result in multiple, low precision estimates of the parameter. This results in a decrease in parameter uncertainty. N_Param_: Total number of fitted parameters, N_Metab_: number of metabolite concentration parameters, N_Nuisance_: number of spectral fitting parameters not of direct interest (e.g., line broadening), N_Model_: number of dynamic model parameters.

For each separation, the estimated amplitude increase was extracted (specified in the GLM as the beta for the rectangular-function stimulation regressor). The RMSE across all repetitions was calculated for the independent and dynamic fits, and the ratio of the uncertainties was calculated (ratio of standard deviations, for both free and linked conditions, Figure 2a&b).

**Figure 2.**
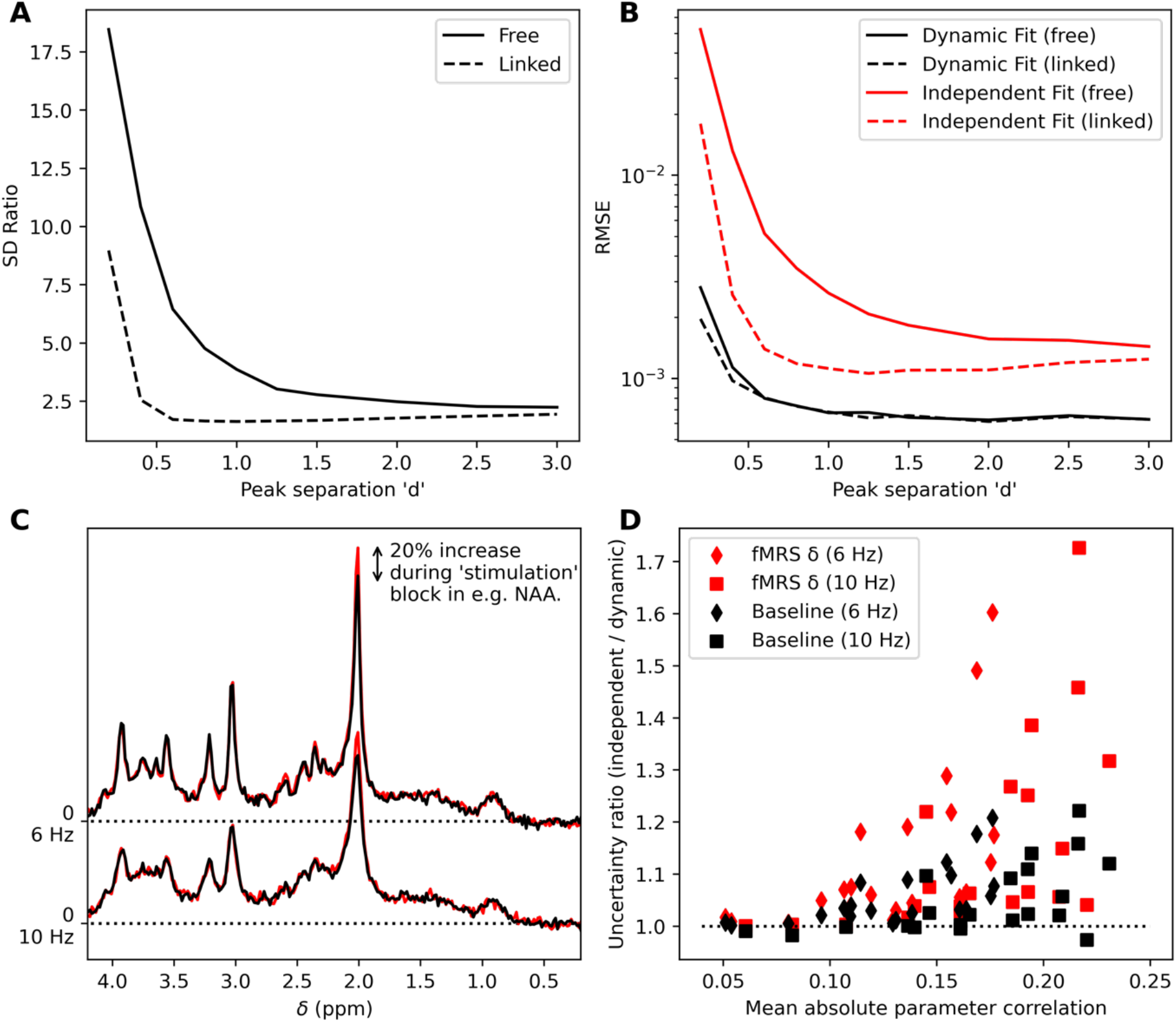
Results of the functional MRS validation. **A** Ratio of Monte Carlo measured standard deviations (independent fitting / dynamic fitting) for the concentration increase as a function of peak separation in the toy two-peak simulation (see Supporting Figure S3). Results for a model with all parameters unlinked “Free” and the standard FSL-MRS fitting model “linked” are given (see §Functional MRS – Simulation). **B** RMSEs for the same simulation. As shown in **A&B** Dynamic fitting reduces uncertainty and overall error. **C** Extension of fMRS validation to realistic 1H-MRS data. Paired data with 20% increases in concentration were simulated for each metabolite (NAA shown) at two linewidths. **D** The uncertainty ratio (ratio of standard deviations, independent fitting / dynamic fitting) for each metabolite’s baseline concentration and increase (delta) is shown as a function of the parameter’s mean correlation with other parameters. A value > 1 indicates that dynamic fitting is decreasing the uncertainty compared to independent fitting.

**Figure 3.**
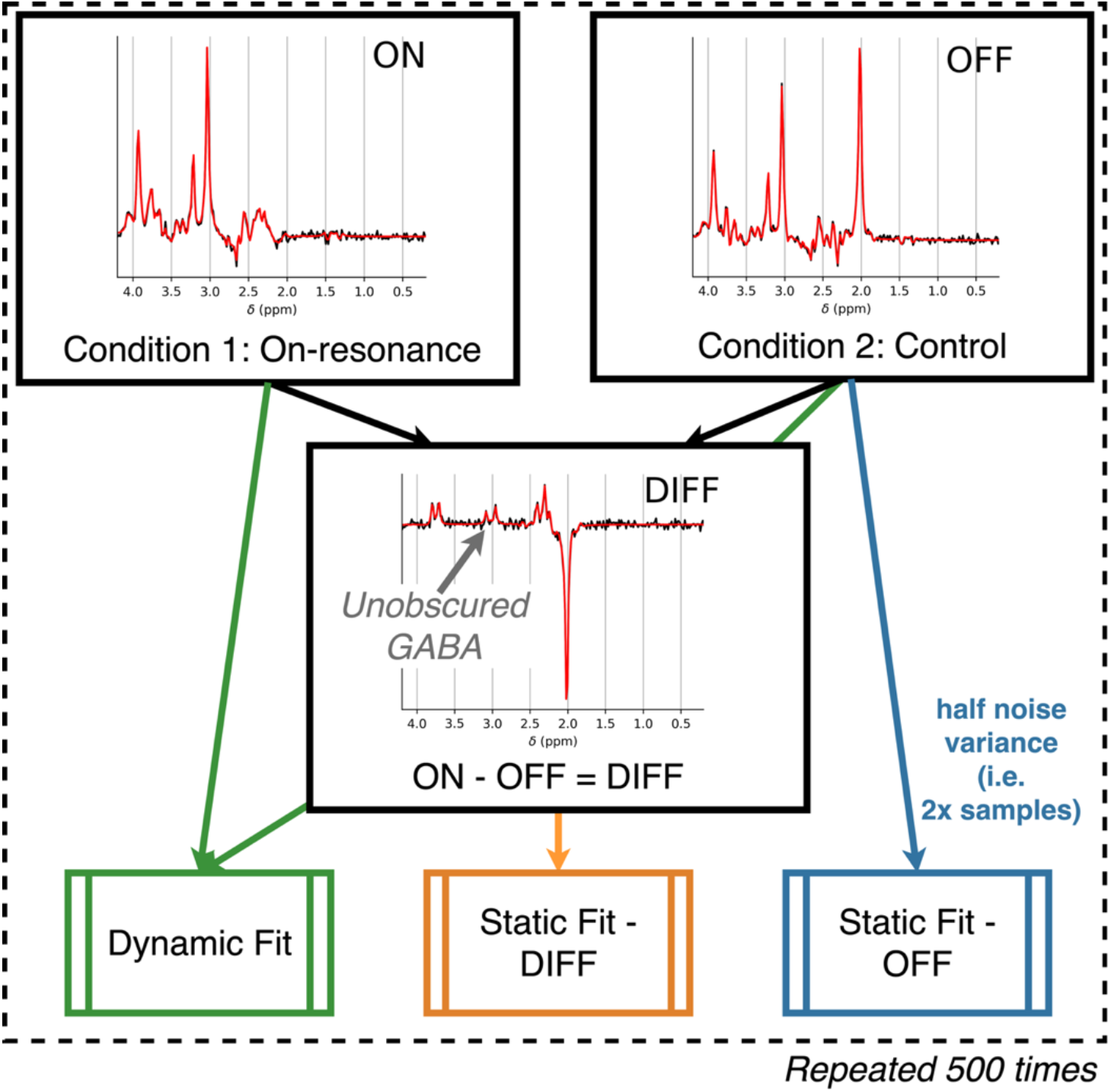
Approach to edited MRS analysis simulation. Simulation is carried out by generating pairs of synthetic MEGA-edited spectra (both the control [OFF] and on-resonance [ON] saturation case), and the corresponding difference spectrum (by subtraction, [DIFF]). The DIFF and OFF spectra are fit using single spectrum fitting, and the ON + OFF spectra are fit using the dynamic approach. The statically fitted OFF spectrum is constructed with half the noise variance to simulate matched time acquisitions. This is repeated 500 times in a Monte Carlo approach for each noise level and line broadening. Spectra are shown with static fitting and have the lowest linewidth (5 Hz) and intermediate noise (noise SD = 144).

The code used to generate and analyse these simulations is contained in the online repository under `./fmrs/1_two_peak_simulation` (git.fmrib.ox.ac.uk/wclarke/fsl-mrs-dynamic-fitting/-/tree/master/fmrs/1_two_peak_simulation).

The second simulation extended the above approach to realistic spectral profiles. In addition, noting that peak separation is a key driver of parameter correlation, and therefore improvement is predicted for the proposed method, the simulation was carried out at two different linewidths (6 Hz and 10 Hz). As such, the same overall approach was taken as the first simulation but implemented with simulated three tesla 1H-MRS spectrum from the brain. For each of 20 metabolites in the spectrum (see supporting information), and for each of the two linewidths, 500 Monte Carlo repetitions were made where one specified metabolite in each simulation increased in amplitude by 20% (example for NAA in Figure 2c) for the central 30 repetitions of 60 total repetitions. All other metabolites were held constant for that case, with the specified metabolite changed for each subsequent case.

Fitting was carried out as for the previous simulation with dynamic fitting implementing a GLM dynamic model using a design matrix with two regressors. No BOLD-like effects on linewidths were simulated.^12^

For each metabolite (and linewidth case) the ratio of independent/dynamic fitting uncertainties (calculated as the standard deviation cross the 500 Monte Carlo repetitions) was calculated for the baseline and stimulation regressor beta. The average concentration parameter correlation was calculated as shown in Supporting Figure S4.

The code used to generate and analyse these simulations is contained in the online repository under `./fmrs/2_fmrs_spectrum_simulation` (git.fmrib.ox.ac.uk/wclarke/fsl-mrs-dynamic-fitting/-/tree/master/fmrs/2_fmrs_spectrum_simulation).

The first (toy two-peak) fMRS simulation shows that in all cases the dynamic fitting approach reduces the uncertainty of the amplitude increase parameters, also showing a lower RMSE. The functional form of the uncertainty ratio as a function of peak separation ‘d’ replicates that found by Tal (Figure 4 in Reference 13). Linking the linewidth and shift parameters, as is done in the default FSL-MRS model, reduces the advantage of dynamic fitting, but retains the same functional form.

**Figure 4.**
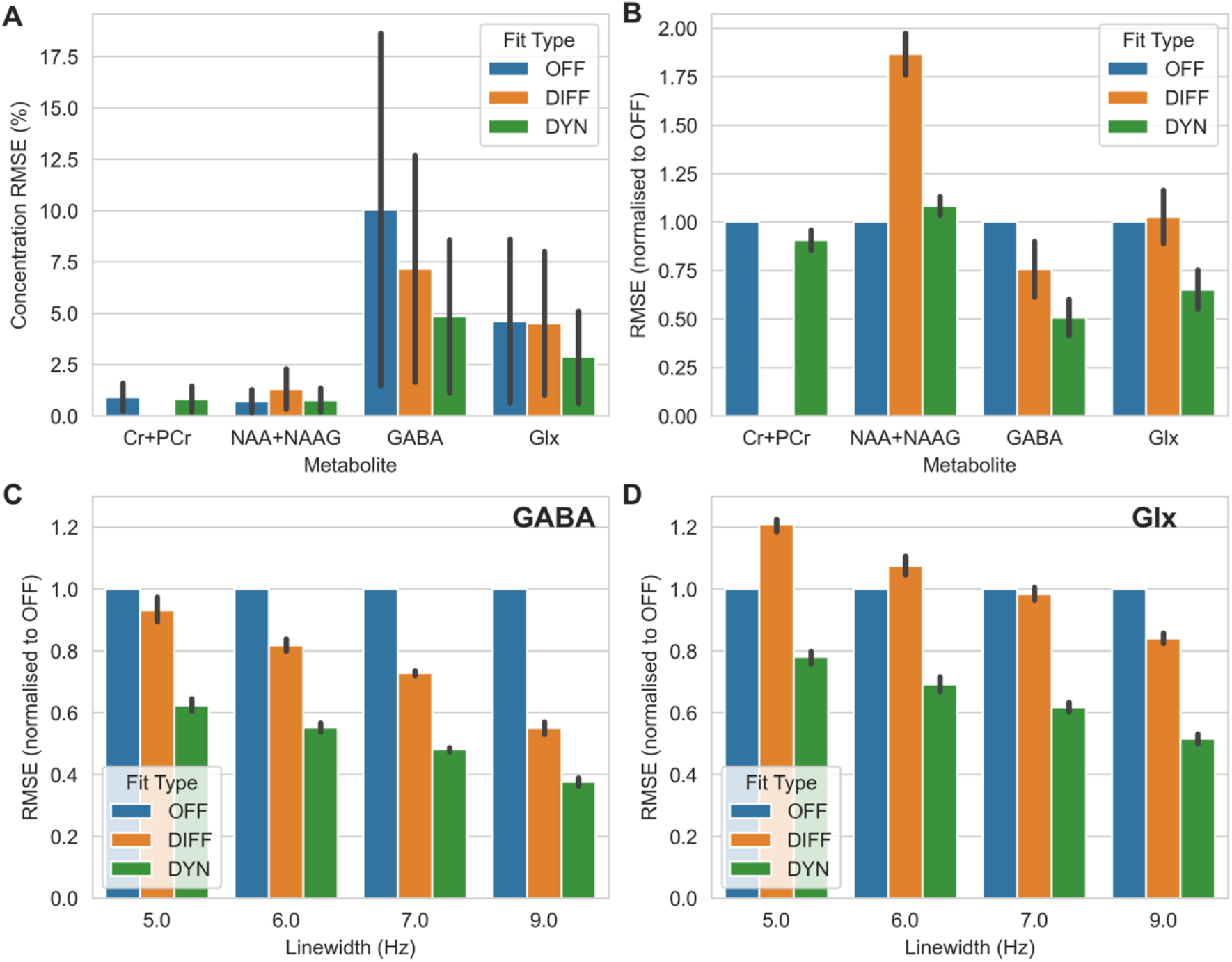
Results of the editing simulation. **A** RMSE (±SD) across all noise levels and linewidths for each examined metabolite, expressed as percentage of the true metabolite concentration. **B** As **A**, but with the results normalised to OFF for each metabolite. **C&D** The effect of linewidth on the relative performance for GABA and Glx (glutamate + glutamine). In all cases, except the measurement of NAA+NAAG, the RMSE is lowest for the dynamic approach.

The second simulated fMRS data, designed to explore data similar to real-world use cases, also demonstrates the advantage of using dynamic fitting over independent fitting for estimating both fMRS amplitude changes and also underlying baseline concentrations. A clear relationship between mean parameter correlation and uncertainty reduction was observed, with wider linewidths giving higher correlations and larger improvements (Figure 2d).

### CS2. Edited-MRS: improved estimation of [GABA]

The second case study uses synthetic single voxel MEGA-PRESS data.^8^ This example is representative of a study that acquires data using the MEGA-PRESS sequence, in absence of an external stimulation paradigm, to measure the concentration of metabolites (e.g. GABA) that are obscured by, or highly correlated with, other metabolite signals. Normally MEGA-PRESS acquires two encoding conditions (ON and OFF), the difference of which (DIFF) contains a simplified spectrum enabling unobscured estimation of GABA. Here, the accuracy and precision on measurements of metabolites (specifically: NAA, creatine, GABA and Glx [glutamate and glutamine]) are compared across three fitting strategies:

1. ‘OFF’ -Control-only acquisition – Taking only the control saturation condition, without a subtraction stage, like an unedited spectrum. All metabolites are visible but many overlap. The spectrum is fitted with a single (unmodified) set of basis spectra.
2. ‘DIFF’ -Forming a difference spectrum – This approach matches the current gold-standard approach. An on-resonance saturation condition is subtracted from a control saturation condition to leave a spectrum containing the differences arising from j-coupling (and direct saturation effects). The difference spectrum is fitted using a modified set of basis spectra.
3. ‘DYN’ -Dynamic fitting of control and on-resonance acquisitions – The proposed approach, control and on-resonance saturation conditions are used in analysis, but no subtraction is performed, and they are analysed together using the proposed simultaneous fitting approach. In this case each spectrum is fit with a relevant set of basis spectra (simulated with control and on-resonance saturation) with additional dynamic constraints. These constraints are equal metabolite concentrations and nuisance parameters (lineshape, shift, phase, baseline, etc.).

In each case the total acquisition time was kept constant, i.e., case one (control-only) data was simulated with half the noise variance. Basis sets were simulated using FSL-MRS’s simulator (`fsl_mrs_sim`), simulations incorporated spatial resolution and fully described RF pulse shapes. The difference basis set was constructed from the subtraction of the OFF from the ON basis set, which respectively simulated editing pulses at 7.5 and 1.9 ppm.

Data was simulated for standard in vivo concentrations for 19 metabolites (specified, with concentrations, in the supporting information), no macromolecules were simulated. Data was simulated with Lorentzian linewidths (FWHM) in four steps from 5 Hz to 9 Hz (representing ‘excellent’ to ‘Acceptable’ linewidths as defined in Reference 22), and eight SNR levels (NAA match-filter SNR of 30 – 330 in 8 steps) that span (and extend beyond) the range observed in vivo. Each condition was simulated 500 times to carry out Monte Carlo sampling of the fitting process. Data was fit using FSL-MRS’s core fitting routine `fit_FSLModel` (parameters specified in supporting information) or the dynamic fitting approach as detailed for fitting case #3. Simulation code for this section is contained in the online repository under `/editing` (git.fmrib.ox.ac.uk/wclarke/fsl-mrs-dynamic-fitting/-/tree/master/editing).

For each metabolite and each fitting condition (#1-3) the root-mean-squared-error (RMSE) was calculated across all Monte Carlo repetitions. RMSE was expressed in both metabolite units (equivalent to mM) or normalised to the time-matched control-only (#1) condition.

The three fitting strategies were simulated and results from four representative metabolites are shown (Figure 4): the frequent targets of MEGA editing, GABA and Glx (glutamate and glutamine combined), a metabolite that appears in all conditions, tNAA (NAA + NAAG), and one which is removed in the differencing process, tCr (creatine + phosphocreatine). Additionally, results are shown for four different linewidths for GABA and Glx (Figure 4: C&D).

For GABA, RMSE was always worst (highest) for OFF (#1), then DIFF (#2) and the lowest was the proposed method DYN (#3). The greatest improvement for DIFF or DYN was seen for widest (worse) linewidths, with DYN achieving an RMSE of 0.39 of the OFF condition with a linewidth of 9 Hz compared to 0.6 for 5 Hz. Across all linewidths DYN achieved a 33% reduction in GABA RMSE compared to DIFF. A similar relationship was seen for Glx, except DIFF was the worst performing fit strategy with narrow linewidths, DYN was always the best.

For tCr and tNAA, DYN produced highly similar results to OFF, both of which substantially outperformed DIFF. No significant variation was observed as a function of SNR or of linewidth for tCr or tNAA (Supporting Figures S5 and S6).

### CS3. fMRS: simulated analysis and group statistics

A full set of simulated visual-stimulation data and analysis scripts has been created for the purpose of demonstrating fMRS analysis using the proposed dynamic fitting approach. The data simulates single-voxel data acquired using block (flashing checkerboard) visual stimulation at 7T, in ten subjects, with a separate stimulation and control condition for each subject. Metabolite concentration changes, inter-subject variance in concentration changes and spectral quality is matched to reported values.^12^ As such, glutamate and lactate were set to increase during stimulation and glucose and aspartate to decrease, on average all other metabolites should be constant. Line narrowing due to the positive BOLD effect was simulated. The input dynamic model uses the canonical BOLD haemodynamic response function to model all changes (metabolite concentrations and line narrowing) during the stimulation period, and is implemented in a design matrix for GLM with four regressors (two stimulation conditions, linear drift, and a constant for modelling baseline concentration, Figure 5B). The simulation implementation is detailed in the supporting information. Group level analysis was conducted using the `fmrs_stats` function from FSL-MRS, implementing a paired t-test design across the stimulation and (no stimulation) control datasets of each subject.

**Figure 5.**
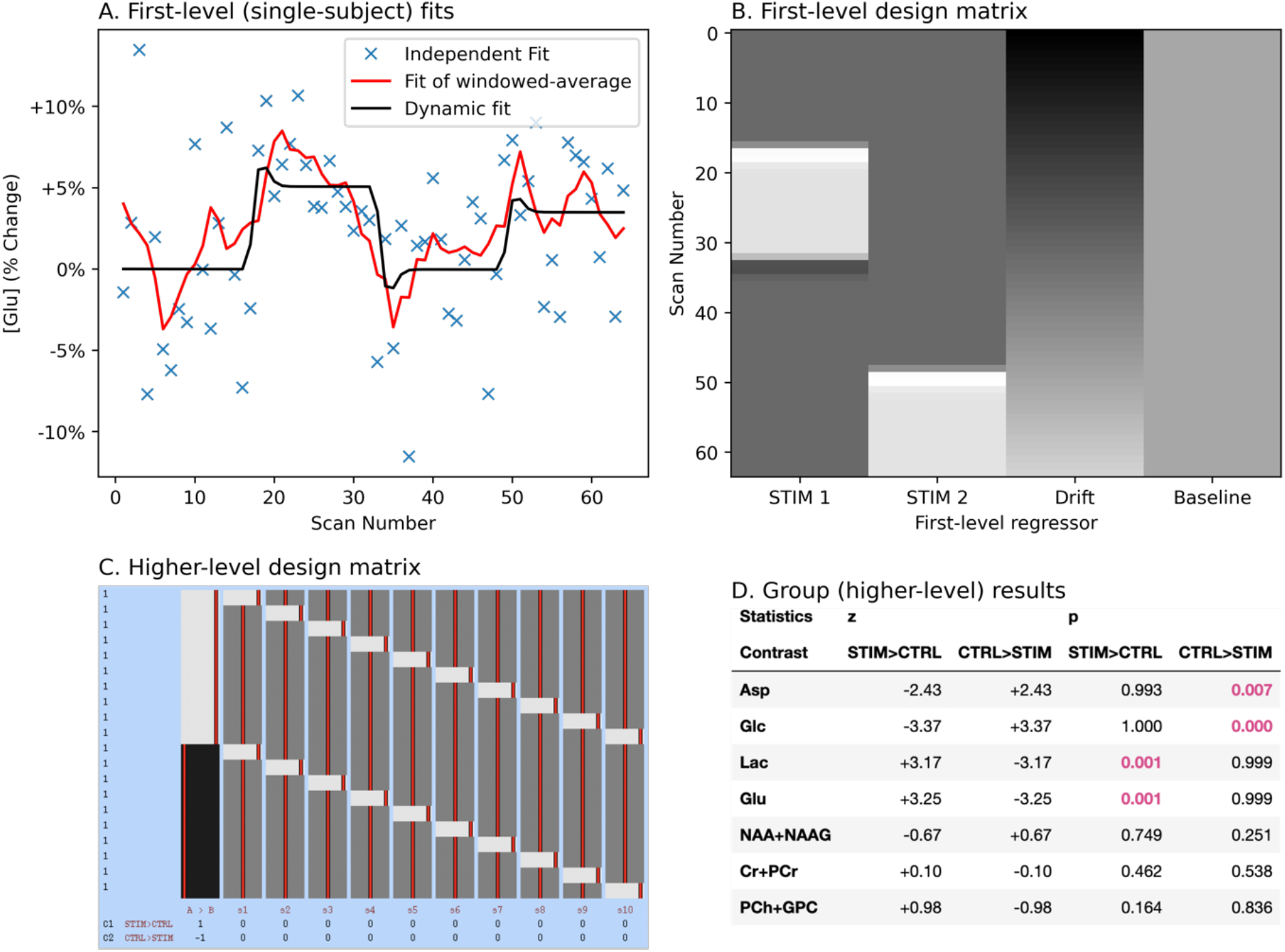
Demonstration of analysis of functional MRS using a GLM at the subject (first-level) and group-level. **A** single subject fit of glutamate. A single subject’s stimulation data is shown for relative glutamate changes. Independent, moving-window temporally smoothed, and GLM/dynamically modelled results are shown. Note that no formal comparison is made to the moving-window averaging method, a formal comparison of dynamic fitting to independent fitting + GLM is made in CS1. **B** Design matrix used to both generate and fit the fMRS data at the first level. There are two stimulation blocks separated by rest blocks, a linear drift regressor, and a constant regressor. **C** The simulated data contained paired stimulation and control (no effect) datasets for ten subjects. The group level analysis used this design matrix (created and displayed using FSL tools) to run a paired t-test. **D** The results as output by FSL-MRS’s fmrs_stats tool. FSL flameo is used to calculate z and p statistics for each first-level contrast at the group level. The tool accurately identifies the metabolites changing in the simulation as significant

This documented demonstration dataset and analysis is hosted separately at github.com/wtclarke/fsl_mrs_fmrs_demo, with a permanent record at Reference 23. In addition to demonstration, this dataset was used to assess the implementation accuracy of the proposed dynamic fitting for fMRS combined with the packaged MRS group-level statistics tool (`fmrs_stats`). To assess the implementation accuracy, the betas for each concentration related regressor was compared to the true simulation input value, per subject and as a group average. In addition to the demonstration code repository (see above), code for this section’s analysis and figure generation is contained in the online repository under `./ fmrs/3_fmrs_demo` (git.fmrib.ox.ac.uk/wclarke/fsl-mrs-dynamic-fitting/-/tree/master/fmrs/3_fmrs_demo).

Implementation accuracy was assessed by comparing GLM betas related to metabolite concentrations (constant and stimulation terms) per subject and at the group level. Figure 6 shows the graphical outputs of group level traces for major metabolites. Lac, Glu, Asp, and Glc were simulated with stimulation-associated concentration changes.

**Figure 6.**
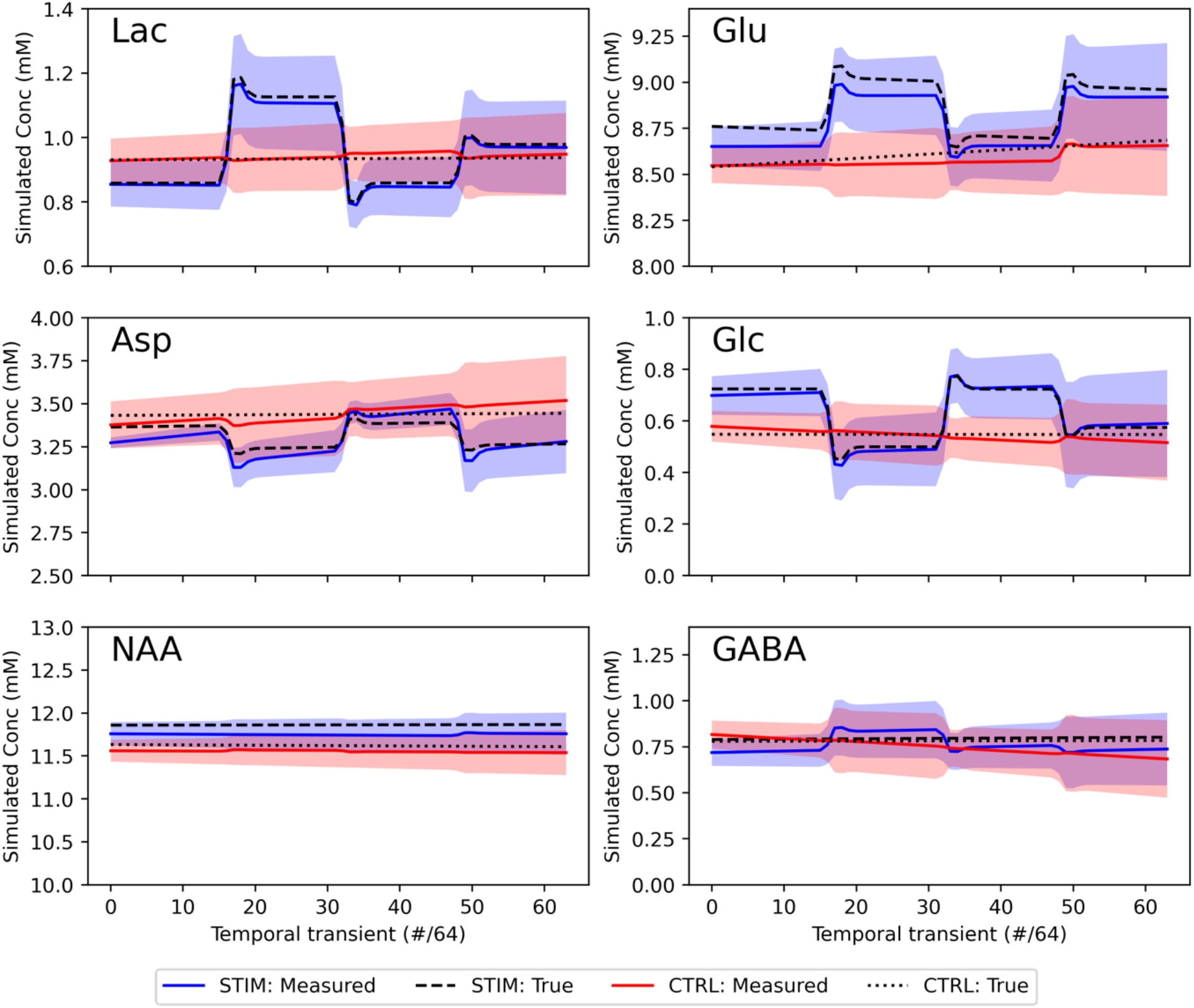
Visual representation of the group-level results from the fMRS demonstration. Stimulation and control (no stimulation) cases are plotted as a function of temporal transient showing group mean and standard deviation. The true values (calculated as mean across subjects) are shown as dashed/dotted lines. All metabolites that had simulated changes (Lac & Glu – positive, Asp & Glc – negative) are shown alongside two with no changes (NAA – high SNR, GABA – low SNR).

**Figure 7.**
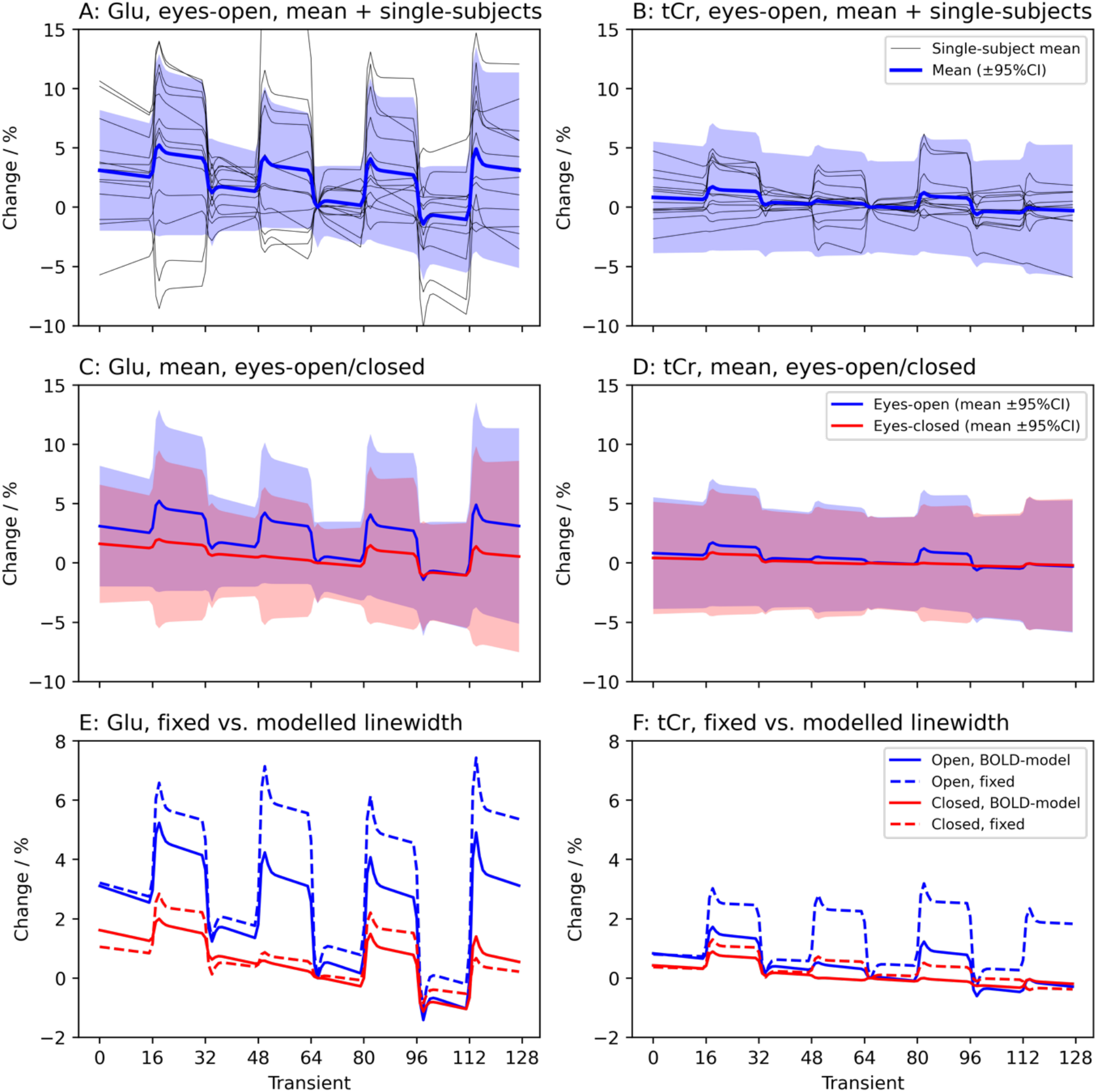
Results of the fMRS in vivo validation for two metabolites: glutamate, which is expected to increase with stimulation (Glu, left) and total creatine which is not expected to change (tCr, right). **A&B** Group-level mean and 95% CI (coloured), and single-subject concentrations expressed as a percentage change relative to the middle time point. **C&D** Comparison of the group level means and 95% CIs for the stimulation (eyes-open) and control (eyes-closed) case. **E&F** Comparison of the results when using a model that incorporates the effect of BOLD on metabolite linewidths, to one with linewidths that are fixed across all timepoints (see supporting information). When fixed spuriously large changes are observed during stimulation, including for non-modulating metabolites. There are much smaller and more variable differences for the control case (where no BOLD effect is expected).

Correlation and Bland-Altman analysis of individual concentration betas is presented in the supporting information (Supporting Figures S7 and S8) and summarised here. Measured betas showed a very high level of correlation at both individual subject level (Pearson’s r = 0.98, bias = 0.4 mM) and group (Pearson’s r = 0.91, bias = 0.02 %).

### CS4. fMRS: In Vivo Confound Mitigation

The proposed dynamic fitting approach was assessed by reanalysing previously published visual stimulation fMRS data.^24^ The original study implements commonly applied analysis approach of carrying out independent spectral fitting on temporally averaged data, and then correlating metabolite time courses (which are themselves further smoothed). The dataset comprises 13 subjects scanned for 8.5 minutes per condition. Two conditions were acquired: a stimulation condition, ‘eyes-open’, with four blocks of flashing checkerboard visual stimulation presented for 64 seconds interleaved with 64 s rest (no stimulation) blocks, and a control condition with ‘eyes-closed’ (no stimulation presented, i.e. constant for 8.5 minutes). Signal was acquired using a sLASER sequence interleaved with 3D EPI, full details are available in the original publication^24^ and are summarised in the supporting information. Note that the interleaved EPI data of the original dataset was not used in this reanalysis. Human data included in this work was collected with informed, written consent, approved by the University of Oxford Research Ethics Committee (MSD-IDREC-C1-2014-146).

Spectral pre-processing was carried out using FSL-MRS’s pre-processing routine, `fsl_mrs_preproc`. The data was analysed using a dynamic fitting approach implementing a GLM model for dynamic analysis. The design matrix was implemented with four stimulation regressors, a linear drift and constant term, generated using the Glover HRF in the Nilearn package.^25,26^ The spectral fitting component used the original study’s basis spectra set (further parameters are listed in the supporting information). Two different approaches to BOLD-induced line narrowing were tested, one where the linewidths were kept fixed across time, and one where the Lorentzian line-broadening was modelled as a GLM (using the same design matrix as metabolite concentrations). This aims to remove the step of applying line broadening to spectra during stimulation events, as carried out in the original study (and others)^12,24^ by multiplying the time-domain data by an exponential filter. Applying line broadening will result in autocorrelation of spectral points and modify noise properties in only the stimulation case.

Group level analysis was carried out using FSL-MRS’s `fmrs_stats` routine implementing a paired t-test design across stimulation and control conditions for each subject (as in CS3).Group results were compared with the original study’s findings.

Code for this section is contained in the online repository under`./fmrs/4_fmrs_invivo_example` (git.fmrib.ox.ac.uk/wclarke/fsl-mrs-dynamic-fitting/-/tree/master/fmrs/3_fmrs_invivo_example).

The original study found significant increases in glutamate, rising approximately 2% over baseline, no other assessed metabolite was found to change during stimulation. In this reanalysis, when BOLD-induced line narrowing was modelled in the GLM the same result was found, with only glutamate showing a statistically significant (at p < 0.05) increase (Supporting Table 1). Glutamate was found to increase 3.1±5.8% on average across all subjects and blocks (p=0.04). A significant decrease in linewidth during the stimulation blocks was found for the eyes-open case (−0.18±0.5 Hz), but not for the eyes-closed case (see Supporting Figure S9). This shows that the line narrowing was successfully modelled without introducing autocorrelation across the signal.

When the BOLD induced narrowing was not modelled in the fixed linewidth model, and nor was it accounted for in processing by line broadening as in the original study (i.e. expected BOLD line narrowing was not accounted for), an increase in glutamate upon stimulation was also found (with increased magnitude, 4.4±5.9%, and significance, p=0.01), but also significant increases in tNAA (0.4±1.0%), tCho (1.9±2.2%), and tCr (1.9±1.1%) were detected. Full statistics are reported in Supporting Table 2.

### CS5. dMRS: Multi-direction Diffusion Encoding

Simulated multi-direction diffusion weighted MRS data was used to demonstrate dynamic fitting enabling new data acquisition approaches for dMRS. By implementing dynamic fitting using a parameterised functional model of direction-dependent diffusion properties, acquisitions that have higher dynamic encoding resolution may be possible, when without dynamic fitting repeated sampling of each encoding would be needed for sufficient SNR. This case study also demonstrates the requirement for adequate fitting initialisation, and how this can be provided using existing tools and the implemented API interface.

Data was simulated (Figure 8a) for three metabolites: NAA, predominantly located neuronally, myo-inositol predominantly located in glia, and creatine, a mix; the simulation was performed for a voxel with two crossing neuronal fibre populations, which is not present for glia. Thus, different simulated metabolites had different orientational diffusion dependence, i.e. different “fibre” orientation distribution function, fODF (simulations values in Supporting Information). NAA parameters were designed to mimic two crossing fibre populations, Ins as a predominantly spherical compartment (mimicking glia), and Cr was implemented as a mixture of the two. Synthetic data was simulated for a diffusion weighted sLASER sequence implementing two diffusion weighting approaches:

**Figure 8.**
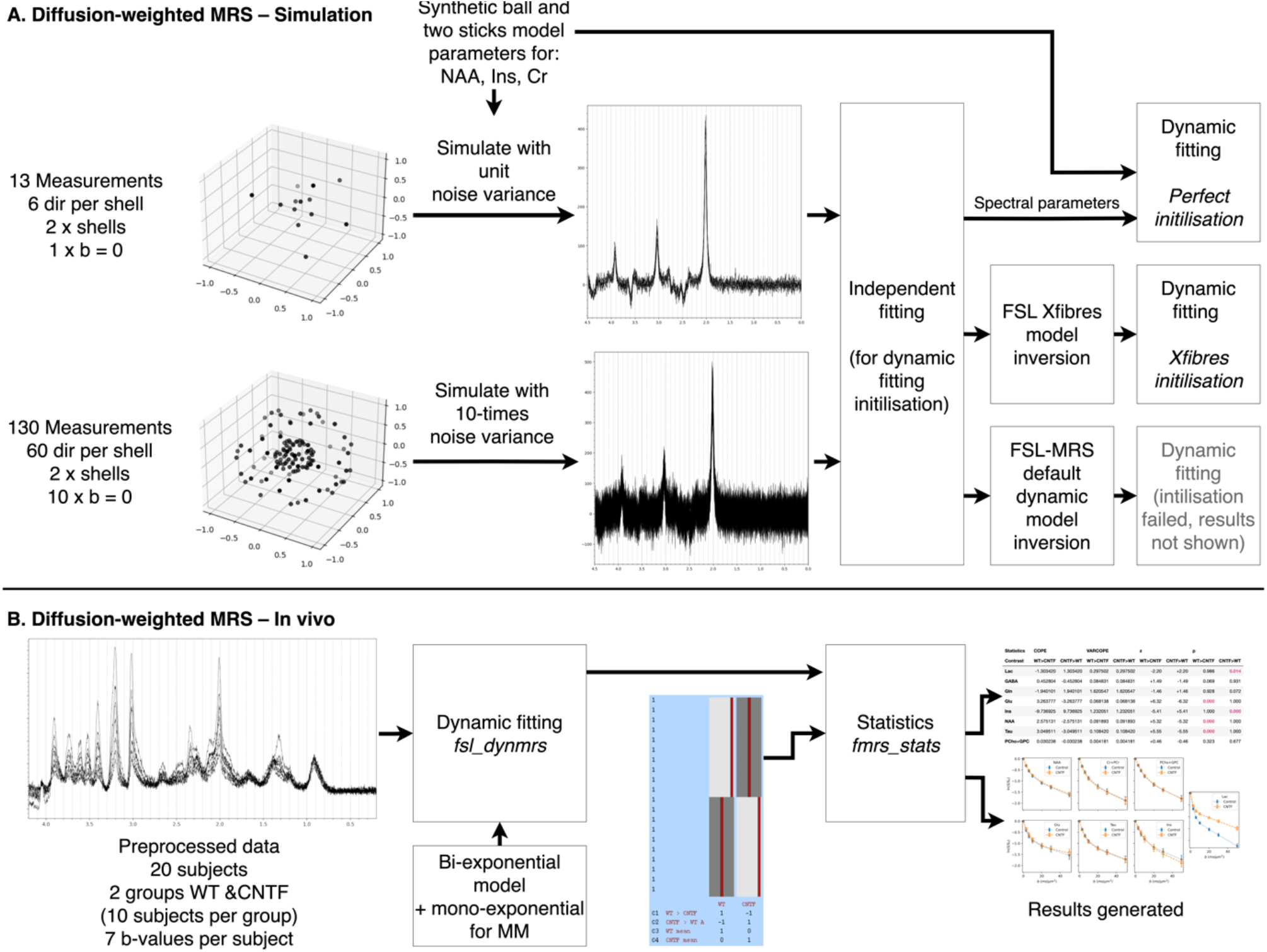
dMRS. **A** Schematic of the simulated analysis of multi-direction data using FSL-MRS’s dynamic approach. Time-matched data with different numbers of diffusion directions was analysed implementing a ball-and-two-sticks model into the spectral-dynamic fitting. Different fitting initialisation approaches were trialled for each case. **B** Previously published multi-b-value dMRS data was reanalysed using spectral-dynamic fitting, implementing a biexponential model (exponential for macromolecules). The group-level analysis using fmrs_stats was qualitatively compared to the published results.

1. Six diffusion directions at two b values (b = 1 and 3 ms/μm^2^), plus b=0, and,
2. Sixty diffusion directions at two b values (b = 1 and 3 ms/μm^2^), plus b=0.

Simulated noise variance was set ten times higher for condition #2, to simulate equal acquisition times. Dataset #2 is what is commonly acquired for water diffusion for modelling crossing fibres (e.g. for tractography), whilst dataset #1 is closer to a dMRS design when repeated measurements are typically needed to increase SNR.

Dynamic fitting was implemented with a two-sticks and a ball diffusion model, which was also used to generate the synthetic data.^27^ The fitting was initialised with one of three approaches:

1. Inversion of the dynamic model using the independently fitted spectra (default in `fsl_dynmrs`),
2. With the ground truth parameters,
3. Using results from the independently fitted spectra passed to FSL’s xfibres routine (which uses a more robust initialisation of the nonlinear fitting tailored to the ball and sticks model).^27^

Quantitative assessment of fitting performance was made using Euclidian distance between ‘stick’ vectors (scaled by ‘fibre’ fraction) estimated to those generated from ground truth parameters. Vector direction was rectified before error calculation.

Code for this section is contained in the online repository under`./dwmrs/1_simulation_dti` (git.fmrib.ox.ac.uk/wclarke/fsl-mrs-dynamic-fitting/-/tree/master/dwmrs/1_simulation_dti).

Estimation of the simulated diffusion parameters (ball and two sticks model in three metabolites) was carried out in four cases, with 6 directions and 60 directions per diffusion shell and using a perfect initialisation strategy (using the ground truth) and using that obtained from the initial independent fit plus FSL’s xfibres routine. Xfibres is the core component of FSL’s Bayesian Estimation of Diffusion Parameters Obtained using Sampling Techniques for crossing fibres (BEDPOSTX), so is designed to estimate the equivalent problem for imaging data of water.^27^ For all metabolites the 60 directions outperformed the 6 directions, and the perfect initialisation outperformed the xfibres approach (Figure 9). The reduction in error between 6 directions to 60 directions (error measured as Euclidian distance between predicted and true fibre vectors) was 23% for the perfect initialisation and 82% for xfibres. Fitting using the default initialisation strategy (inversion of the model in FSL-MRS) failed in all cases due to a complex loss landscape resulting in multiple local minima, and the results are not shown.

**Figure 9.**
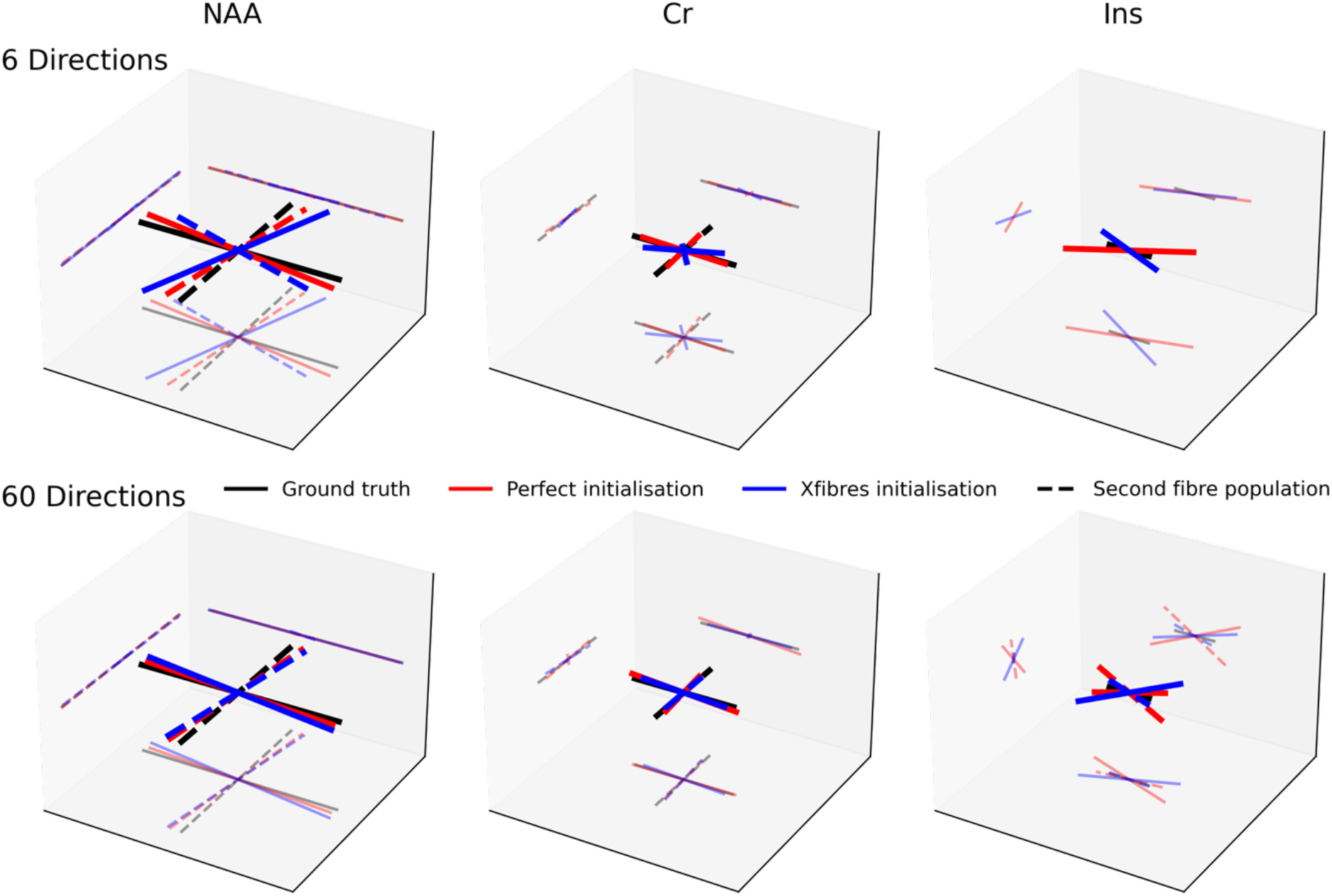
Results of fitting the ball and two-sticks model to simulated multi-direction diffusion data. This is a demonstration of the ability of the tool to simultaneously fit more complex diffusion models and spectral information. However, a good initialisation point (provided by FSL’s xfibres tool) is required. Here simulated data with more diffusion directions (but correspondingly lower spectral SNR) provides a better estimate of fibre directions than lower numbers of directions, which is required for stable spectral fits when no information is shared. The xfibres initialised fit, achievable on real data, is compared with an artificial perfect initialisation approach (which requires the ground truth) and the ground truth. Each metabolite simulates a different cellular compartmentalisation and therefore has a different ground truth.

**Figure 10.**
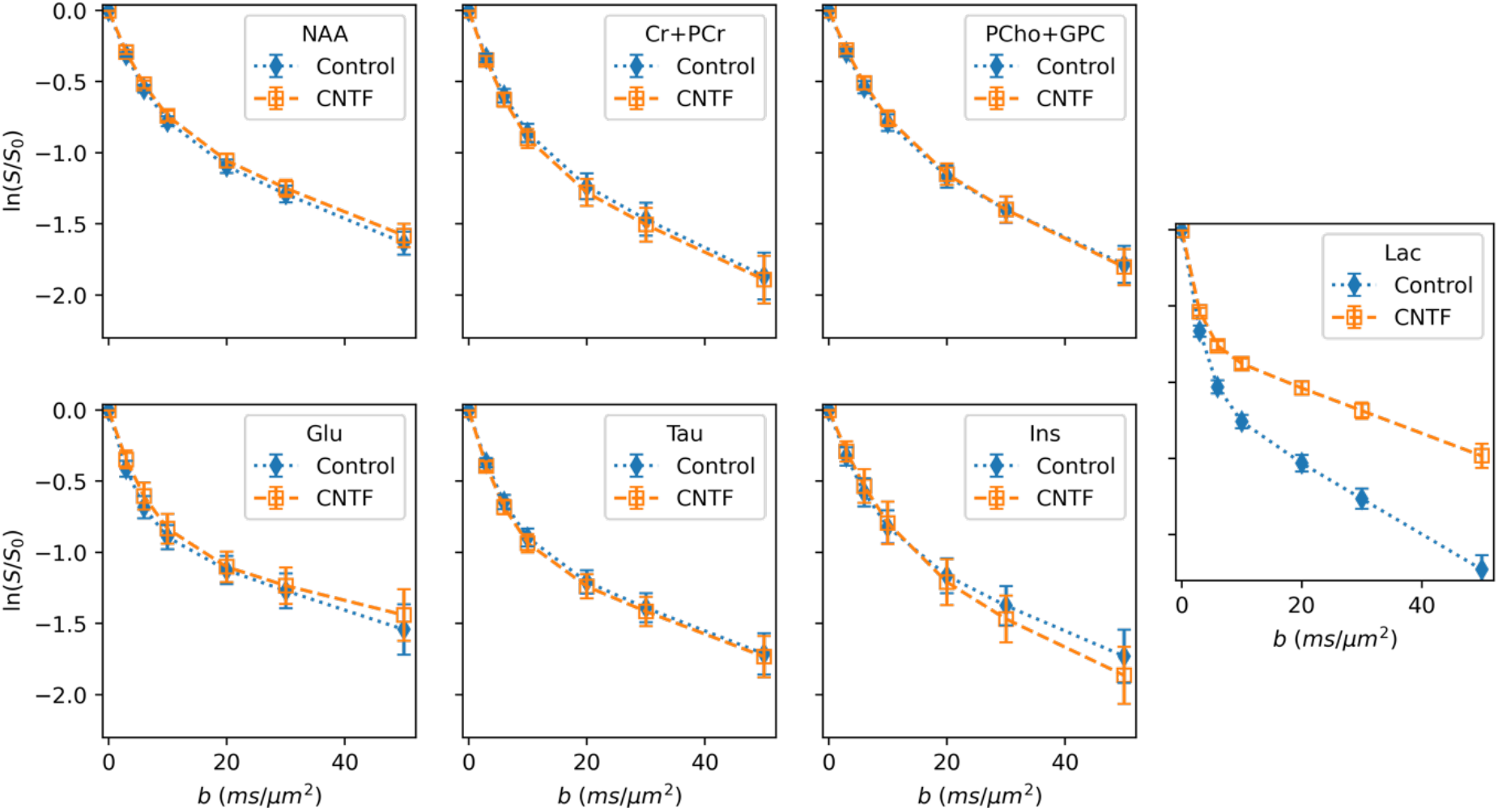
Results of the in vivo dMRS validation. The concentrations of key metabolites as fitted using the proposed dynamic fitting approach. The two cohorts are shown separately. The results closely match the magnitude and direction of the original publication’s results, which found significant differences in myo-inositol (Ins) and lactate (Lac) diffusion properties. Changes in overall metabolite concentrations were also found (see Supporting Table 3), again matching the original publication. This analysis used a bi-exponential representation to fit the dynamic process.

### CS6. dMRS: in vivo validation

Dynamic fitting of diffusion-weighted MRS was demonstrated on a previously published dataset of a mouse model of “pure” astrocyte reactivity induced by injecting cytokine ciliary neurotrophic factor (CNTF).^28^ Two groups of ten mice (control and CNTF), were scanned using a diffusion-weighted STE-LASER sequence,^29^ acquiring data at seven b-values ranging from 0.02 to 50 ms/μm^2^. Acquisition details are summarised in the supporting information.

Processed data was provided by the original study’s authors. Dynamic fitting implemented a bi-exponential model for metabolite concentrations, and a mono-exponential model for macromolecules. An exponential-plus-offset model was used for polynomial baseline terms, this accounts for decreasing baseline ‘roll’ arising from unmodelled residual water signal, which decreases b-values increase. Without this baseline model either the baseline must be estimated independently for each b-value, increasing variability from a purely data-driven component of fitting, or a fixed value must be used, increasing bias. The spectral basis contained 20 metabolites, including empirically measured macromolecules (see supporting information).

Comparison was made to the original published results which implemented independent fitting using LCModel,^17^ and applied complex models of diffusion in randomly oriented cylinders.^6^

Code for this section is contained in the online repository under `./dwmrs/2_invivo` (git.fmrib.ox.ac.uk/wclarke/fsl-mrs-dynamic-fitting/-/tree/master/dwmrs/2_invivo).

The results of the proposed fitting approach accurately recapitulate the results of the original study,^28^ both looking at microscopic properties as measured using diffusion weighting and overall metabolite concentrations (supporting information). Here we find statistically significant differences in myo-inositol and lactate diffusion, and statistically significant differences in metabolite concentrations for Lac, Glu, Ins, NAA, Tau. The original publication found the same changes, apart from lactate concentration differences which did not reach statistical significance in the original, but we note that Bonferroni correction was applied in the original.

## Discussion & Conclusion

As demonstrated here, our universal dynamic MRS fitting tool allows simultaneous analysis of multiple, linked spectra. The tool implements a novel framework capable of handling several common types of dynamic MRS and incorporates methods for doing higher level statistics. We used the tool to replicate published results from fMRS and dMRS studies. We also numerically replicated theoretical results showing improvements in fitting uncertainty when using dynamic fitting approaches. This will help mitigate the low SNR inherent in dynamic spectroscopic approaches, improving target parameter precision. It will also enable greater dynamic resolution by permitting finer dynamic sampling despite the concurrent SNR reduction per shot.

In this work we have demonstrated that on in vivo data the tool successfully replicates the results of published work from the original data, which used independent fitting approaches. On simulated data, where a ground truth is available, we demonstrated that the tool accurately estimates dynamic model parameters for diffusion weighted, functional, and edited spectroscopy experiments. Further, it was used to demonstrate an improvement in parameter estimation precision over independent fitting of edited spectroscopy. In addition, this work provides demonstration uses, and practical examples. Documentation has been developed beside the software tool and is available at *fsl-mrs*.*com*.

The tool also integrates into a larger neuroimaging toolbox, FSL, and uses standardised data formats, compatible with neuroimaging (NIfTI and NIfTI-MRS).^30,31^ By doing this, the tool will allow integration with existing neuroimaging methods (as demonstrated by the *fmrs_stats* tool) and use of well validated MRI approaches, for example physiological noise regression in fMRS and mixed-effects group level modelling as shown in our dynamic fMRS results in this paper. To this point, in the toolbox, and in the examples shown, we have integrated existing models and software tools developed for analogous MRI techniques (fMRS/fMRI, dMRS/dMRI).

Although our toolbox is designed to handle any type of dynamic spectroscopy, some practical limitations remain. Currently the tool does not implement models which require dynamic parameters linked to more than one metabolite. For example, models where two or more metabolites interact dynamically. This limits applications to e.g., tracer experiments, such as used in hyperpolarised carbon-13 spectroscopy, where the dynamic modelling requires this level of flexibility. This limitation is however only a matter of implementation and will be addressed in future versions.

Finally, this work does not address an important caveat, which is identifying suitable dynamic models for linking across spectra. For example, in our fMRS analyses, we have approximated the response of metabolite concentrations to sensory stimulation using the canonical BOLD fMRI haemodynamic response function. While the BOLD response is well characterised in fMRI, it is not yet so in fMRS. Dynamic fitting using a suboptimal model may bias the results in ways that must be quantified in future studies. Similarly, there are many possible diffusion models already arising from dMRI, but these require adaptation for spectroscopy. Furthermore, there are myriad choices for the user to make during pre-processing, choice of fitting algorithm and model description: where the normal ‘static’ spectral fitting parameters are extended by parameters describing the dynamic model. Changing many of these parameters can change the results, especially given the low SNR measurements we are fitting. This work does not start to address these choices, but the tool and framework presented provide a rigorous analysis and statistical platform to do model discovery and selection.

## Acknowledgements

WTC is funded by Wellcome [225924/Z/22/Z]. SJ is funded by Wellcome [221933/Z/20/Z and 215573/Z/19/Z]. The Wellcome Centre for Integrative Neuroimaging is supported by core funding from the Wellcome Trust [203139/Z/16/Z and 203139/A/16/Z]. IBI is funded by a Royal Society Dorothy Hodgkin Fellowship (DHF\R1\201141).

This research was funded in whole, or in part, by the Wellcome Trust [Grant numbers 225924/Z/22/Z, 203139/Z/16/Z and 203139/A/16/Z]. For the purpose of open access, the author has applied a CC BY public copyright licence to any Author Accepted Manuscript version arising from this submission.

We thank Clémence Ligneul, Julien Valette, and Betina Ip for providing the in vivo data for this paper.

## Data Availability Statement

The code and data used in this work are available online.

The underlying fitting code is available as part of FSL-MRS (version used 2.1.0, git hash a2dd6c3820c2311289027f2fb41a0d5c431f3917), this is available from fsl-mrs.com, https://git.fmrib.ox.ac.uk/fsl/fsl_mrs/, and a permanent record is available at https://doi.org/10.5281/zenodo.5910341.

The source code specifically for this work is available from https://git.fmrib.ox.ac.uk/wclarke/fsl-mrs-dynamic-fitting (specific git hash c52d6021f3bb64dc89daa80344dcc76c3fa74c2c), with a permanent record at https://doi.org/10.5281/zenodo.7956321. The original, unprocessed data for the in vivo fMRS and dMRS sections is available from https://doi.org/10.5281/zenodo.7950984.

The specific fMRS fitting demonstration code is available from https://github.com/wtclarke/fsl_mrs_fmrs_demo (git hash a1e475ce18fa601a70e5e8b4041217f521f4787a) with a permanent record at https://doi.org/10.5281/zenodo.7951648.

