## Supporting Information for "Universal Dynamic Fitting of Magnetic Resonance Spectroscopy"

### Universal Dynamic Fitting of Magnetic Resonance Spectroscopy – Supporting Information

#### Fitting Method and Initialisation

Fitting initialisation is either performed automatically, by inverting the dynamic model using independently fitted concentrations arising from each time-point spectrum, or by manually provided initial parameters. The independent fits to each time-point spectrum (for initialisation), are themselves initialised by fitting the mean spectrum to mitigate low SNR.

Fitting optimisation is performed using the Truncated Newton algorithm as implemented in Scipy (default),[1,2] or using a Metropolis-Hastings Markov chain Monte Carlo algorithm, as specified by the user.

#### Configuration files

The Python syntax configuration file defines the behaviour of the spectral-dynamic fitting process. The configuration file is briefly described in the main manuscript but is expanded on here with examples. Full configuration files for each section in the methods & results can be found in the accompanying code repository.

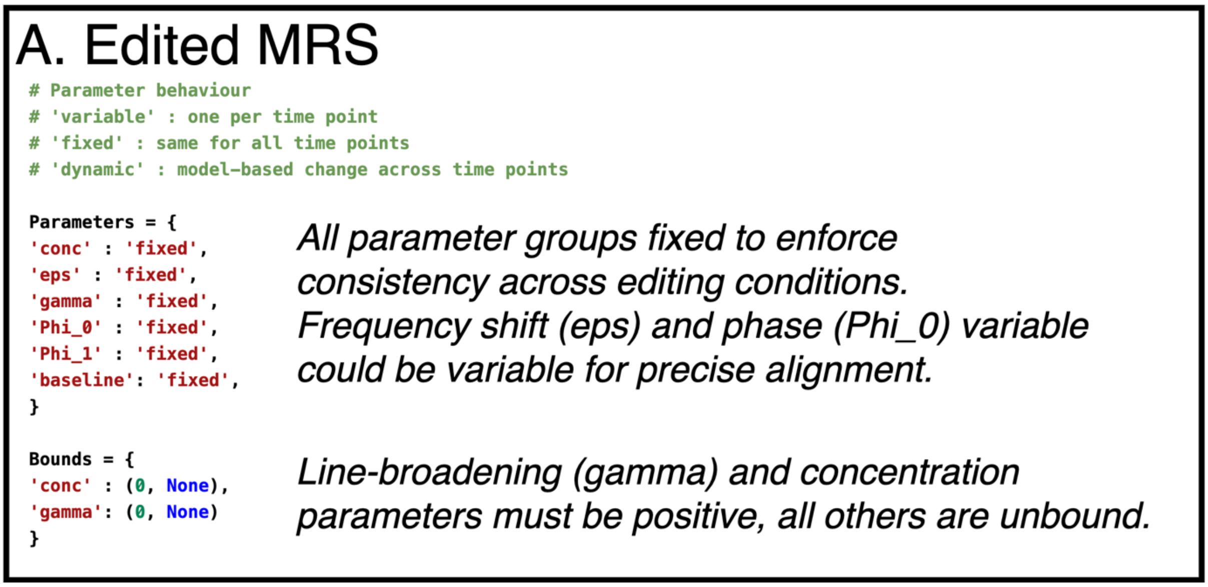


*Supporting Figure 1a. Example configuration file for edited spectroscopy.*

#### B. fMRS

```
from numpy import dot

Parameters = {
'conc' : {'dynamic': 'model_glm', 'params': ['beta_STIM0', 'beta_STIM1', 'beta_LIN', 'beta_CONST']},
'gamma' : 'fixed',
'sigma' : {'dynamic': 'model_glm', 'params': ['beta_STIM0', 'beta_STIM1', 'beta_LIN', 'beta_CONST']},
'eps' : 'fixed',
'baseline' : 'fixed',
'Phi_0' : 'fixed',
'Phi_1' : 'fixed'}
```

*GLM used to model concentrations and Gaussian line-broadening (sigma). All other parameters are fixed to a single value.*

```
Bounds = {
'gamma': (0, None),
'beta_CONST': (0, None)}
```

*Lorentzian (gamma) line-broadening and GLM constant term must be positive.*

```
# Dynamic models
def model_glm(p, t):
return dot(t, p)
```

*Model is dot-product of GLM design matrix and time variable.*

```
# Dynamic model gradients
def model_glm_grad(p, t):
return t.T
```

*Gradient is therefore the time variable.*

*Supporting Figure 1b. Example configuration file for functional spectroscopy fit with a GLM approach. The design matrix (not shown) contains four regressors: two stimulation terms, a linear drift term, and a constant term.*

#### C. dwMRS

```
Parameters = {
'conc' : {
   'MM'    : {'dynamic': 'model_exp',   'params': ['c_amp', 'c_mono_adc']},
   'other' : {'dynamic': 'model_biexp', 'params': ['c_amp', 'c_adc_slow', 'c_adc_fast', 'c_frac_slow']}
   },
'baseline' : {'dynamic': 'model_exp_offset', 'params': ['b_amp', 'b_adc', 'b_off']}
}
```

*A bi-exponential model  is defined for the metabolite concentrations, except the macromolecules which are mono-exponential. An exponential (+ offset) model is used for the baseline.*

```
Bounds = {
'c_amp' : (0, None),
'c_mono_adc' : (0, None),
'c_adc_slow' : (0, .1),
'c_adc_fast' : (.1, 4),
'c_frac_slow' : (0, 1),
'gamma' : (0, None),
'sigma' : (0, None),
'b_amp' : (None, None),
'b_adc' : (1E-5, 3),
}
```

*Bounds are used to impose a consistent ordering between fast and slow ADCs, and enforce a fraction between 0 and 1.*

*Baseline amplitude (b_amp) can be negative unlike concentration. Limit on decay speeds are imposed.*

```
# Dynamic models
from numpy import exp
from numpy import asarray
from numpy import ones_like

# Mono-exponential
def model_exp(p, t):
# p = [amp,adc]
return p[0]*exp(-p[1]*t)
```

*All models (exponential, bi-exponential, exponential + offset) are defined here.*

```
# Mono-exponential model with offset
def model_exp_offset(p, t):
# p = [amp,adc,off]
return p[2]+p[0]*exp(-p[1]*t)

# Bi-exponential model
def model_biexp(p, t):
# p = [amp,adc1,adc2,frac]
return p[0]*(p[3]*exp(-p[1]*t)+(1-p[3])*exp(-p[2]*t))

# Gradients
# For each of the models defined above, specify the gradient
def model_exp_grad(p,t):
e1 = exp(-p[1]*t)
g0 = e1
g1 = -t*p[0]*e1
return asarray([g0,g1], dtype=object)
```

*Analytical expressions are provided for the function gradients Defined with the model function name + "_grad".*

*Bi-exponential and exponential+offset gradient not shown.*

*Supporting Figure 1c. Example configuration file for multi b-value* diffusion-weighted *spectroscopy. Different metabolites are fit using different models (mono-exponential for macromolecules, bi-exponential for all others). An exponential-plus-offset is used for the polynomial baseline terms, which change as residual water is supressed by diffusion sensitisation. All other parameter sets are fixed. Gradients (not all shown) are precalculated for these simple analytical models.*

### CS1. Functional MRS: replication, and extension of Tal

#### Methods

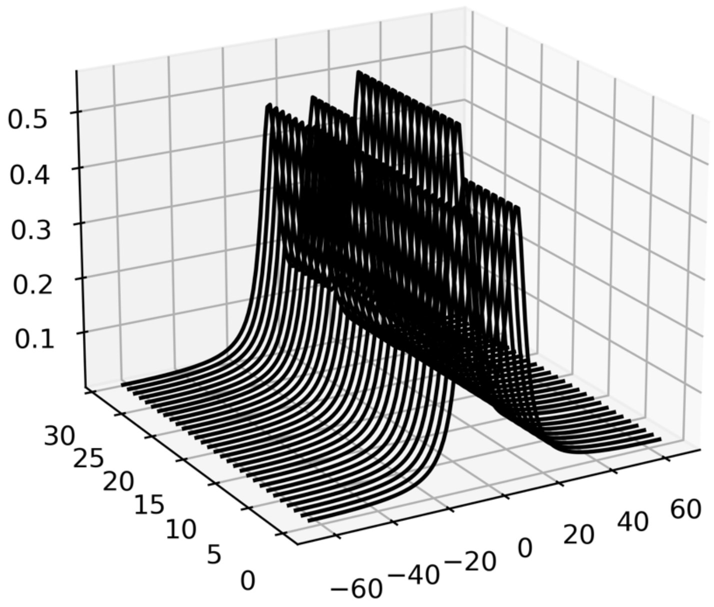


*Supporting Figure 2. fMRS model used in the first "two-peak" simulation. 64 spectral transients are created, each containing two peaks (every other shown). During the central 32 transients one peak increases in amplitude by a fractional amount $\delta$. The other peak has a constant amplitude in all transients. The separation of the peaks can be varied (see Supporting Figure 5).*

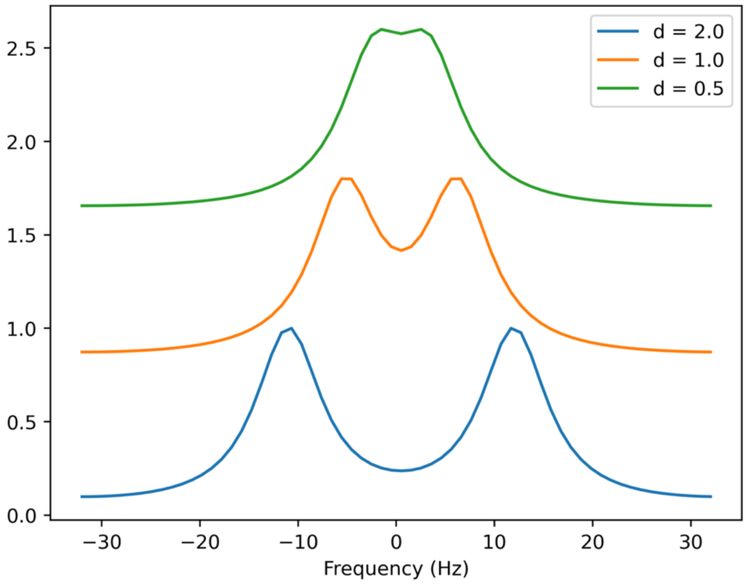


*Supporting Figure 3. Peak separation examples for the first "two-peak" simulation. Each spectral transient contains two Lorentzian peaks, with equal, 8 Hz linewidth. The separation of the peaks can be controlled using a factor d, where the total separation in hertz is* $8\sqrt{2}\mathrm{d}$.

For the simulation of fMRS synthetic spectra were created with 20 metabolites: Ala, Asc, Asp, Cr, GABA, Glc, Gln, Glu, Gly, GPC, GSH, Ins, Lac, NAA, NAAG, PCh, PCr, PE, Scyllo, Tau. Simulated concentrations are listed in the table below. One metabolite, iterating through all 20 in each simulation, underwent a 20% increase in concentration during stimulation.

| **Metabolite** | **Concentration (mM)** | **Metabolite** | **Concentration (mM)** |
|---|---:|---|---:|
| Alanine (Ala) | 0.60 | Glutathione (GSH) | 1.20 |
| Ascorbate (Asc) | 1.20 | Myo-Inositol (Ins) | 7.72 |
| Aspartate (Asp) | 2.40 | Lactate (Lac) | 0.60 |
| Creatine (Cr) | 4.87 | N-acetyl aspartate (NAA) | 13.80 |
| GABA | 3.50 | N-acetyl aspartate glutamate (NAAG) | 1.20 |
| Glucose (Glc) | 1.20 | Phosphorylcholine (PCh) | 0.85 |
| Glutamine (Gln) | 3.37 | Phosphocreatine (PCr) | 4.87 |
| Glutamate (Glu) | 12.41 | Phosphorylethanolamine (PE) | 1.80 |
| Glycine (Gly) | 1.20 | scyllo-Inositol (Scyllo) | 0.30 |
| Glycerophosphorylcholine (GPC) | 0.74 | Taurine (Tau) | 1.80 |

During simulation the synthetic data was created and fit using the following controllable parameters: Lorentzian lineshapes, 0$^{th}$-order polynomial baseline, single metabolite group (all metabolite have same line-broadening and no relative frequency shifts allowed), optimisation region of 0.2 to 4.2 ppm.

#### Results

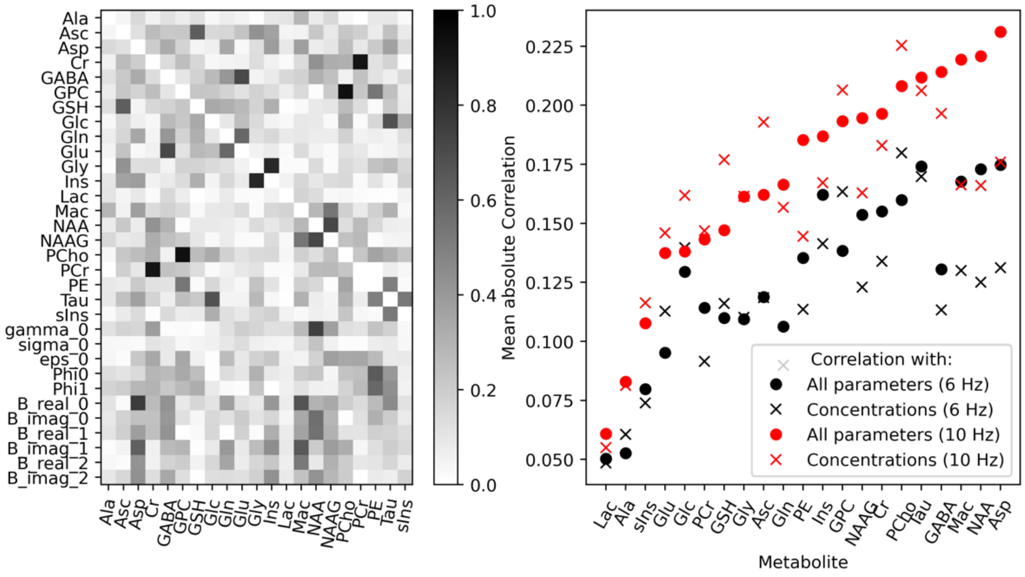


*Supporting Figure 4. Left: Mean parameter correlation matrix arising from repeatedly fitting a synthetic spectrum used in the second fMRS simulation (50 repetitions used). This was repeated for both linewidth conditions (6 Hz and 10 Hz) and used to calculate the mean correlations. Right: The sorted mean correlations between each metabolite concentration parameter and either all other concentrations or all parameters (i.e. concentrations & nuisance parameters). The mean all parameter correlation was used in the main analysis (Figure 6D).*

### CS2. Edited-MRS: improved estimation of [GABA]

#### Methods

For the simulation of edited spectroscopy synthetic spectra were created with 19 metabolites: Ala, Asc, Asp, Cr, GABA, Glc, Gln, Glu, GPC, GSH, Ins, Lac, NAA, NAAG, PCh, PCr, PE, Scyllo, Tau. Simulated concentrations are listed in the table below:

| **Metabolite** | **Concentration (mM)** | **Metabolite** | **Concentration (mM)** |
|---|---:|---|---:|
| Alanine (Ala) | 0.60 | Myo-Inositol (Ins) | 7.72 |
| Ascorbate (Asc) | 1.20 | Lactate (Lac) | 0.60 |
| Aspartate (Asp) | 2.40 | N-acetyl aspartate (NAA) | 13.80 |
| Creatine (Cr) | 4.87 | N-acetyl aspartate glutamate (NAAG) | 1.20 |
| GABA | 3.50 | Phosphorylcholine (PCh) | 0.85 |
| Glucose (Glc) | 1.20 | Phosphocreatine (PCr) | 4.87 |
| Glutamine (Gln) | 3.37 | Phosphorylethanolamine (PE) | 1.80 |
| Glutamate (Glu) | 12.41 | scyllo-Inositol (Scyllo) | 0.30 |
| Glycerophosphorylcholine (GPC) | 0.74 | Taurine (Tau) | 1.80 |
| Glutathione (GSH) | 1.20 | | |

During analysis simulation the synthetic data was fit using the following controllable parameters: Lorentzian lineshapes, $0^{th}$-order polynomial baseline, single metabolite group (all metabolites have the same line-broadening and no relative frequency shifts allowed), optimisation region of 0.2 to 4.2 ppm.

#### Results

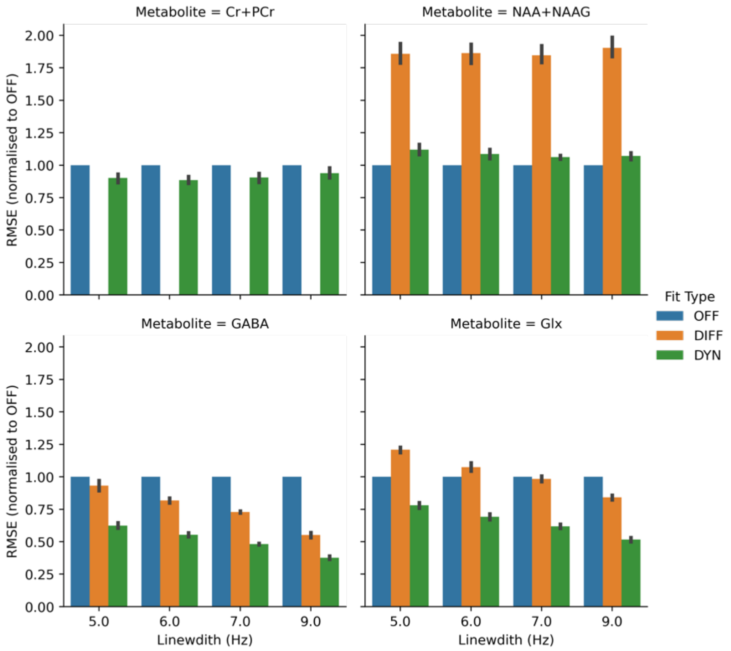


*Supporting Figure 5. Effect of linewidth on relative performance of different edited MRS fitting strategies. Performance is measured using RMSE normalised to OFF and expressed as a mean ± SD across all Monte Carlo repetitions and noise levels. Note the DIFF method cannot measure total creatine (Cr+PCr), so is not reported. Note that the lower two panels are the same as Figure 5 panels C & D.*

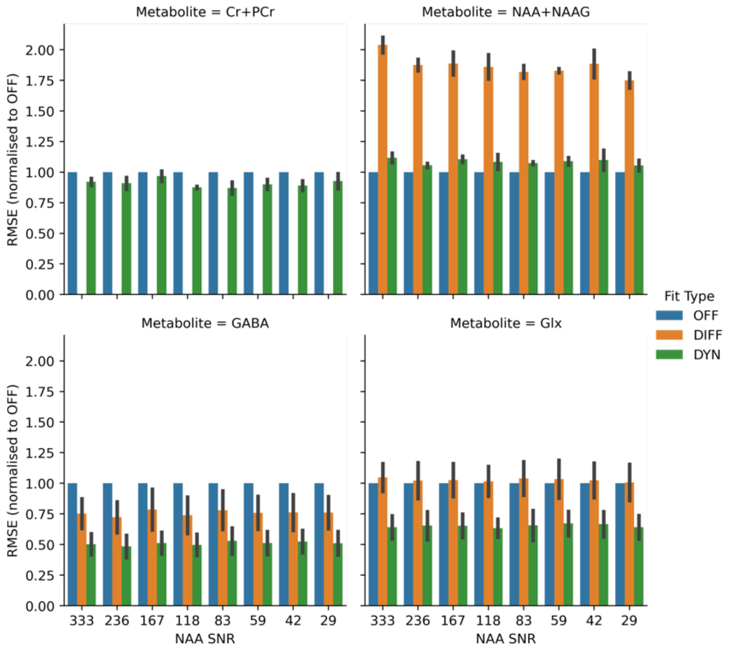


*Supporting Figure 6. Effect of noise level (measured as NAA SNR) on relative performance of different edited MRS fitting strategies. Performance is measured using RMSE normalised to OFF and expressed as a mean ± SD across all Monte Carlo repetitions and linewidths. Note the DIFF method cannot measure total creatine (Cr+PCr), so is not reported.*

### CS3. fMRS: simulated analysis and group statistics

#### Methods – Simulation detail

The data in this case study comprises two simulated fMRS datasets (with and without stimulation) from each of 10 subjects. Each dataset comprises 64 spectra with two blocks of stimulation and two blocks of rest, in the pattern (REST-STIM-REST-STIM).

Four metabolite concentrations change during the stimulation period, with the changes being drawn from population distributions as reported in Reference 3. Specifically, Glutamate, Lactate, Glucose, Aspartate following the these changes:

- Glutamate = 4±1.3% rise during STIM
- Lactate = 25±23% rise during STIM
- Glucose = -16±18% drop during STIM
- Aspartate = -5±4% drop during STIM

The changes were modelled using Nilearn's *make_first_level_design_matrix* function, implementing the 'Glover' HRF model with stimulation blocks 16 transients long (TR=4 s) starting at the 16$^{th}$ and 48$^{th}$ transient. A 1$^{st}$ order polynomial drift and constant term was included. All other metabolites are constant w.r.t STIM. Random linear drift term of up to ±5% applied to each metabolite.

A gaussian linewidth narrowing of 0.5 Hz during STIM following the response of the HRF is applied to all metabolites. Control data has no concentration changes during STIM, nor linewidth changes, but did include a random linear drift.

The simulated data had complex noise added, which has a per-subject standard deviation drawn from a distribution generated from the SNR measured from local fMRS studies.

For precise implementation details see the file *generate_data.ipynb* in the online repository: github.com/wtclarke/fsl_mrs_fmrs_demo.

#### Results

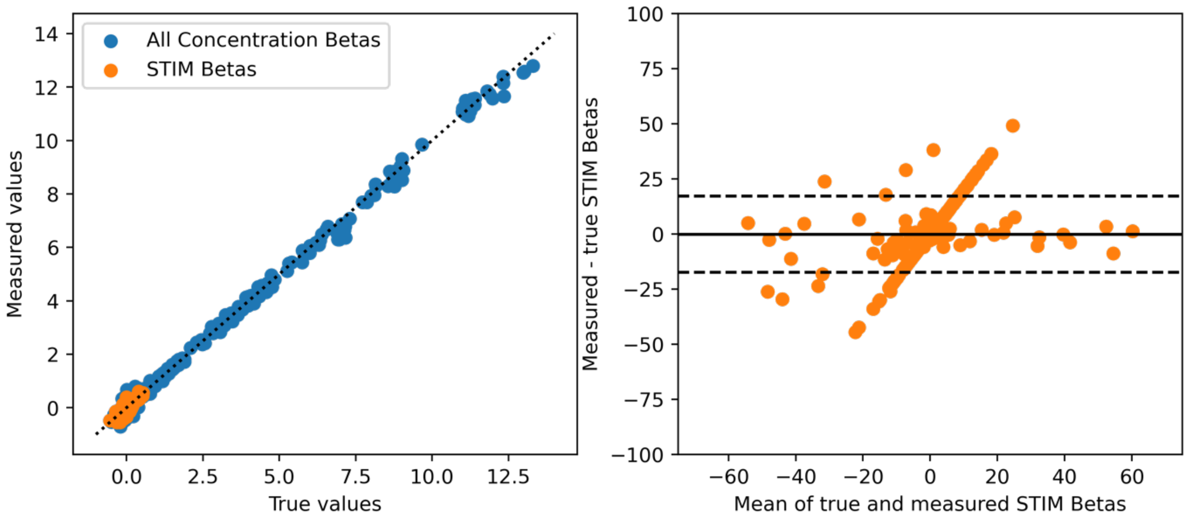


*Supporting Figure 7. Correlation (left) and Bland-Altman plot (right) of the measured vs true first-level (i.e. per subject) beta values in the fMRS demo. All betas (stimulation, constant*

*and drift) are shown in blue, while just the stimulation betas are shown in orange. The diagonal line of parameters in Bland-Altman analysis arises from 0-valued betas which are estimated to have small, non-zero values.*

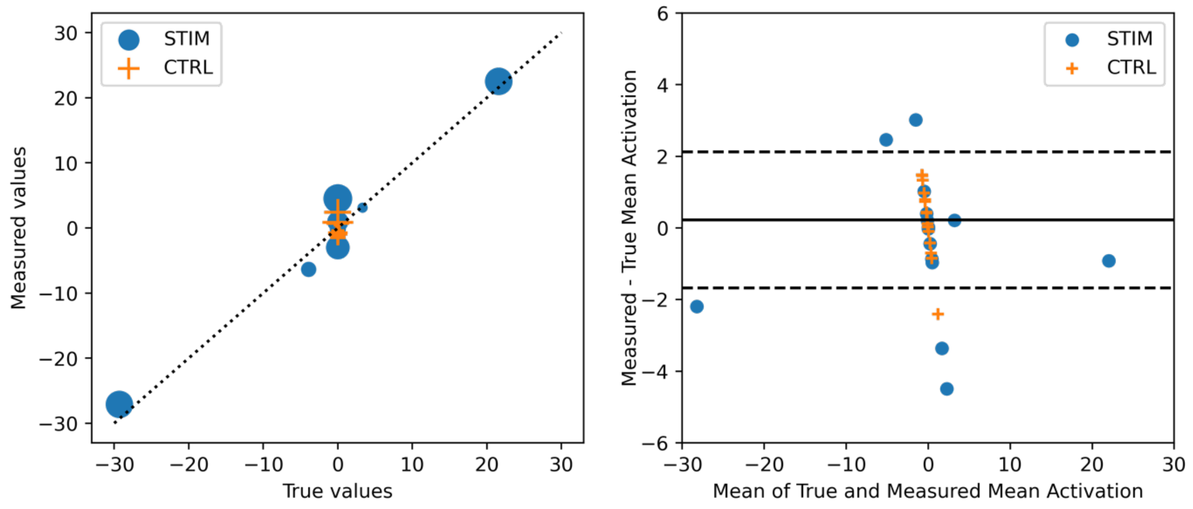


*Supporting Figure 8. Correlation (left) and Bland-Altman plot (right) of the measured vs true group-level 'mean activation' beta values in the fMRS demo. Stimulation condition betas) are shown in blue, while the control condition betas are shown in orange. The diagonal line of parameters in Bland-Altman analysis arises from 0-valued betas which are estimated to have small, non-zero values. Mean activation is the mean value of the two stimulation betas.*

#### CS4. fMRS: In Vivo Confound Mitigation

##### Methods

Data was supplied by the authors of the original study, Reference 4. Briefly, data from eighteen volunteers (9 females, mean age 28.71 ± 5.62 years) was collected using a 7 T whole body MR-scanner (Siemens, Erlangen) with a Nova Medical head coil (single transmit, 32 receive channels). After the original authors' QC process, 13 subjects were included (for more details, see Reference 4), these were used in this work. A 2×2×2 $cm^3$ MRS voxel was positioned in the occipital lobe, centred along the midline and the calcarine sulcus. Data was acquired using a joint fMRI-MRS sequence.[5] Here only the MRS data was used. MRS data were acquired with a semi-LASER sequence (TE=36 ms, TR=4 s), with VAPOR water suppression and outer volume suppression.[6,7] Data was pre-processed using `fsl_mrs_preproc` which applies phase & frequency correction, eddy current correction (using the water reference), repetition averaging, residual water removal (HLSVD algorithm), extra FID point truncation (see `fmrs/4_fmrs_invivo_example/0b_preprocess.py` in the online repository for details).

Fitting of the in vivo fMRS data used a basis set provided by the original study authors. It contained 21 metabolites: Ala, Asc, Asp, Cr, GABA, Glc, Gln, Glu, Gly, GPC, GSH, Ins, Lac, NAA, NAAG, PCho, PCr, PE, Scyllo, Tau, and empirically measured macromolecules. Note that in the original study a slightly modified basis set which split the singlet and multiplet components of NAA and the aspartyl, acetyl and glutamate moieties of NAAG was used.

Fitting used Voigt lineshapes, a 1st-order polynomial baseline, two metabolite groups (all metabolites apart from the empirically measured macromolecules have same line-broadening and no relative frequency shifts allowed), optimisation region of 0.2 to 4.2 ppm.

#### Results

| Statistics | COPE | | VARCOPE | | z | | p | |
|---|---|---|---|---|---|---|---|---|
| Contrast | STIM>CTRL | CTRL>STIM | STIM>CTRL | CTRL>STIM | STIM>CTRL | CTRL>STIM | STIM>CTRL | CTRL>STIM |
| Ala | 2.39e-03 | -2.39e-03 | 2.23e-05 | 2.23e-05 | 0.49 | -0.49 | 0.31 | 0.69 |
| Asc | 6.98e-03 | -6.98e-03 | 2.49e-05 | 2.49e-05 | 1.32 | -1.32 | 0.09 | 0.91 |
| Asp | 7.91e-03 | -7.91e-03 | 5.24e-05 | 5.24e-05 | 1.04 | -1.04 | 0.15 | 0.85 |
| GABA | -3.53e-03 | 3.53e-03 | 5.38e-05 | 5.38e-05 | -0.47 | 0.47 | 0.68 | 0.32 |
| GSH | -2.05e-03 | 2.05e-03 | 3.78e-06 | 3.78e-06 | -1.01 | 1.01 | 0.84 | 0.16 |
| Glc | -1.94e-03 | 1.94e-03 | 1.23e-05 | 1.23e-05 | -0.54 | 0.54 | 0.71 | 0.29 |
| Gln | 4.98e-03 | -4.98e-03 | 3.19e-05 | 3.19e-05 | 0.85 | -0.85 | 0.20 | 0.80 |
| Glu | 7.93e-03 | -7.93e-03 | 1.74e-05 | 1.74e-05 | 1.74 | -1.74 | 0.04 | 0.96 |
| Gly | -1.18e-03 | 1.18e-03 | 2.89e-05 | 2.89e-05 | -0.21 | 0.21 | 0.58 | 0.42 |
| Ins | 1.64e-03 | -1.64e-03 | 6.99e-06 | 6.99e-06 | 0.60 | -0.60 | 0.27 | 0.73 |
| Lac | 1.92e-03 | -1.92e-03 | 1.37e-05 | 1.37e-05 | 0.50 | -0.50 | 0.31 | 0.69 |
| PE | 5.32e-03 | -5.32e-03 | 1.63e-05 | 1.63e-05 | 1.25 | -1.25 | 0.11 | 0.89 |
| Scyllo | -2.35e-04 | 2.35e-04 | 5.35e-07 | 5.35e-07 | -0.31 | 0.31 | 0.62 | 0.38 |
| Tau | -7.03e-04 | 7.03e-04 | 7.44e-06 | 7.44e-06 | -0.25 | 0.25 | 0.60 | 0.40 |
| mm | -7.74e-05 | 7.74e-05 | 3.59e-09 | 3.59e-09 | -1.22 | 1.22 | 0.89 | 0.11 |
| NAA+NAAG | 1.44e-03 | -1.44e-03 | 2.55e-06 | 2.55e-06 | 0.87 | -0.87 | 0.19 | 0.81 |
| Cr+PCr | 2.23e-03 | -2.23e-03 | 1.88e-06 | 1.88e-06 | 1.52 | -1.52 | 0.06 | 0.94 |
| PCho+GPC | 5.02e-04 | -5.02e-04 | 3.25e-07 | 3.25e-07 | 0.85 | -0.85 | 0.20 | 0.80 |
| Glu+Gln | 1.33e-02 | -1.33e-02 | 1.64e-05 | 1.64e-05 | 2.72 | -2.72 | 0.00 | 1.00 |

*Supporting Table 1. Statistical group-level results of the paired t-test run on the in vivo fMRS data, using the GLM driven linewidth first-level model. Note the spurious mean activations of the metabolites typified by strong singlet resonances (tNAA: NAA+NAAG, tCr: Cr+PCr, tCho: PCho+GPC) have disappeared (compared to ST2), leaving only significance driven by changes in glutamate (Glu).*

| Statistics | COPE | | VARCOPE | | z | | p | |
|---|---|---|---|---|---|---|---|---|
| **Contrast** | **STIM>CTRL** | **CTRL>STIM** | **STIM>CTRL** | **CTRL>STIM** | **STIM>CTRL** | **CTRL>STIM** | **STIM>CTRL** | **CTRL>STIM** |
| **Ala** | 1.56e-03 | -1.56e-03 | 2.29e-05 | 2.29e-05 | 0.32 | -0.32 | 0.37 | 0.63 |
| **Asc** | 8.37e-03 | -8.37e-03 | 2.23e-05 | 2.23e-05 | 1.64 | -1.64 | 0.05 | 0.95 |
| **Asp** | 1.67e-03 | -1.67e-03 | 4.43e-05 | 4.43e-05 | 0.25 | -0.25 | 0.40 | 0.60 |
| **GABA** | -1.14e-02 | 1.14e-02 | 6.34e-05 | 6.34e-05 | -1.34 | 1.34 | 0.91 | 0.09 |
| **GSH** | -8.69e-04 | 8.69e-04 | 3.79e-06 | 3.79e-06 | -0.44 | 0.44 | 0.67 | 0.33 |
| **Glc** | -4.34e-03 | 4.34e-03 | 1.29e-05 | 1.29e-05 | -1.15 | 1.15 | 0.87 | 0.13 |
| **Gln** | 4.68e-04 | -4.68e-04 | 2.18e-05 | 2.18e-05 | 0.10 | -0.10 | 0.46 | 0.54 |
| **Glu** | 1.21e-02 | -1.21e-02 | 2.15e-05 | 2.15e-05 | 2.28 | -2.28 | 0.01 | 0.99 |
| **Gly** | -1.78e-03 | 1.78e-03 | 3.05e-05 | 3.05e-05 | -0.32 | 0.32 | 0.62 | 0.38 |
| **Ins** | 3.65e-03 | -3.65e-03 | 7.02e-06 | 7.02e-06 | 1.30 | -1.30 | 0.10 | 0.90 |
| **Lac** | 1.87e-03 | -1.87e-03 | 1.36e-05 | 1.36e-05 | 0.49 | -0.49 | 0.31 | 0.69 |
| **PE** | -7.31e-04 | 7.31e-04 | 1.64e-05 | 1.64e-05 | -0.18 | 0.18 | 0.57 | 0.43 |
| **Scyllo** | -1.12e-04 | 1.12e-04 | 5.35e-07 | 5.35e-07 | -0.15 | 0.15 | 0.56 | 0.44 |
| **Tau** | 1.99e-04 | -1.99e-04 | 6.97e-06 | 6.97e-06 | 0.07 | -0.07 | 0.47 | 0.53 |
| **mm** | -5.83e-05 | 5.83e-05 | 3.28e-09 | 3.28e-09 | -0.98 | 0.98 | 0.84 | 0.16 |
| **NAA+NAAG** | 6.30e-03 | -6.30e-03 | 2.55e-06 | 2.55e-06 | 3.10 | -3.10 | 0.00 | 1.00 |
| **Cr+PCr** | 7.41e-03 | -7.41e-03 | 2.36e-06 | 2.36e-06 | 3.53 | -3.53 | 0.00 | 1.00 |
| **PCho+GPC** | 1.50e-03 | -1.50e-03 | 3.25e-07 | 3.25e-07 | 2.30 | -2.30 | 0.01 | 0.99 |
| **Glu+Gln** | 1.24e-02 | -1.24e-02 | 1.52e-05 | 1.52e-05 | 2.66 | -2.66 | 0.00 | 1.00 |

*Supporting Table 2. Statistical group-level results of the paired t-test run on the in vivo fMRS data, using the fixed linewidth first-level model. Note the spurious mean activations of the metabolites typified by strong singlet resonances (tNAA: NAA+NAAG, tCr: Cr+PCr, tCho: PCho+GPC).*

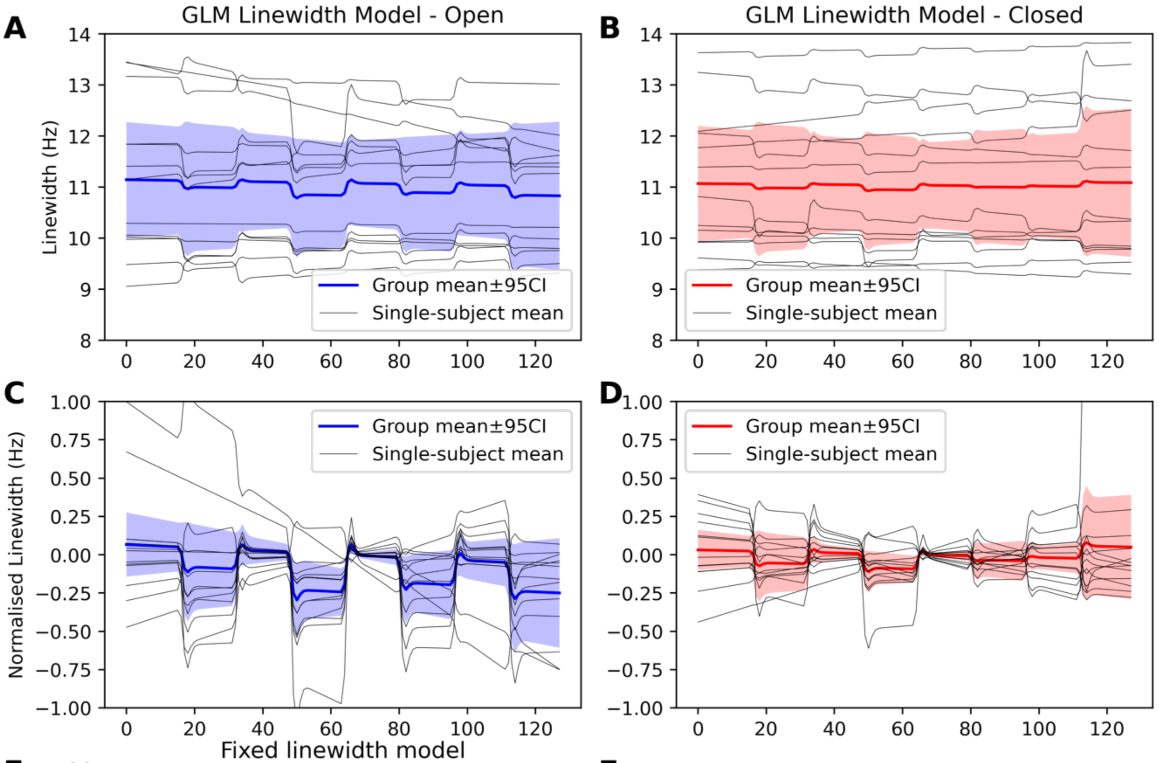


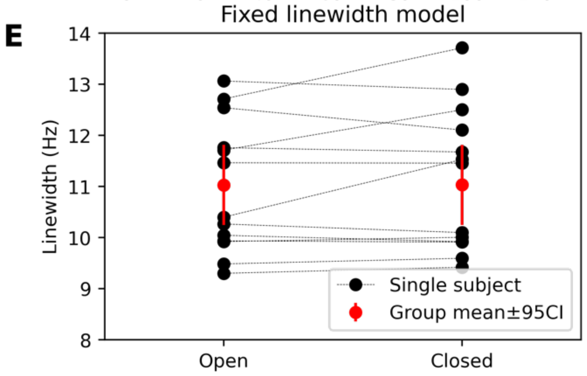


F

| | ΔLW (Hz) | std (Hz) | z stat | p |
|---|---|---|---|---|
| **Closed** | -0.028 | 0.027 | -0.977 | 0.164 |
| **Open** | -0.179 | 0.052 | -2.832 | 0.002 |

*Supporting Figure 9. Behaviour of linewidth in in vivo fMRS.* ***A&B*** *show the estimated linewidth, per-subject and for the group, when modelled using the GLM approach, for the stimulus (open) condition (****A****) and the control (closed) condition (****B****).* ***C&D*** *Show the same but with the constant term removed, so only changes in linewidth are seen. IN the open case a clear decrease in linewidth is seen during the stimulus blocks, which isn't consistently seen on the control case. The statistical analysis (****F****) shows that only in the open condition was there a statistically significant change in linewidth, with a mean decrease of -0.18±0.5 Hz.* ***E*** *The fixed linewidth model showed overall similar linewidths but no change in linewidth was found between the open and closed conditions.*

### CS5. dMRS: Multi-direction Diffusion Encoding

Diffusion model simulation parameters for CS5 are given in the following table. NAA parameters were designed to mimic two crossing fibre populations. Ins as a predominantly spherical compartment. Cr was implemented as a mixture of the two.

| Parameter | Cr | Ins | NAA |
|---|---|---|---|
| s0 | 4.9 | 7.7 | 14 |
| d | 0.00019 | 0.00019 | 0.0002 |
| th1 | 1.6 | 1.6 | 1.6 |
| ph1 | 0 | 0 | 0 |
| f1 | 0.25 | 0.1 | 0.5 |
| th2 | 1.6 | 0 | 1.6 |
| ph2 | 1.6 | 0 | 1.6 |
| f2 | 0.25 | 0 | 0.5 |

### CS6. dMRS: in vivo validation

#### Methods

Data was supplied by the authors of Reference 8. Data was acquired on mice in a 11.7 T Bruker horizontal scanner (maximum gradient strength $G_{max}$ = 752 mT/m) with a transmit-receive quadrature surface cryoprobe (Bruker, Ettlingen, Germany). Animals from two cohorts were scanned. The cohorts were wild type (WT) and those which underwent Cytokine Ciliary Neurotrophic Factor injection (CNTF) to induce astrocytic hypertrophy without neuronal death or microglial activation. Animals were scanned anesthetized, held at constant temperature, and fixed with bite and ear bars. The voxel was in the striatum and 6.5 mm x 3 mm x 2.8 mm (56 $mm^3$). The sequence was a STE-LASER sequence with TE = 33.4 ms, diffusion gradient duration of 3 ms, and VAPOR water suppression.[9] Seven diffusion weightings were (b = 0.02, 3.02, 6, 10, 20, 30 and 50 ms/$\mu m^2$). TR = 2000 ms, 128 repeats per b value (one gradient direction for all repeats). Scan-to-scan phase correction was performed before averaging across repeats. Eddy current correction was achieved using a co-acquired water reference. For more details see Reference 8.

Fitting of the in vivo dwMRS data used a basis set provided by the original study authors. It contained 20 metabolites: Ace, Ala, Asp, Cr, GABA, Glc, Gln, Glu, Gly, GPC, GSH, Ins, Lac, NAA, NAAG, PCho, PCr ,PE ,Tau, and empirically measured macromolecules (measured from the same cohort). Fitting used Voigt lineshapes, a $1^{st}$-order polynomial baseline, two metabolite groups (all metabolites apart from the empirically measured macromolecules have same line-broadening and no relative frequency shifts allowed), optimisation region of 0.2 to 4.2 ppm.

#### Results

In addition to the diffusion parameter results reported in Figure 10 of the main manuscript, both the FSL-MRS dynamic analysis and the original analysis found gross changes in metabolite concentrations. These are reported here, with lactate, glutamate, myo-inositol, NAA and taurine showing statistically significant differences. The original publication found the same changes, less lactate which fell below statistical significance.

| Statistics | COPE | | VARCOPE | | z | | p | |
|---|---|---|---|---|---|---|---|---|
| Contrast | WT>CNTF | CNTF>WT | WT>CNTF | CNTF>WT | WT>CNTF | CNTF>WT | WT>CNTF | CNTF>WT |
| **Lac** | -1.303420 | 1.303420 | 0.297502 | 0.297502 | -2.20 | +2.20 | 0.986 | 0.014 |
| **GABA** | 0.452804 | -0.452804 | 0.084831 | 0.084831 | +1.49 | -1.49 | 0.069 | 0.931 |
| **Gln** | -1.940101 | 1.940101 | 1.620547 | 1.620547 | -1.46 | +1.46 | 0.928 | 0.072 |
| **Glu** | 3.263777 | -3.263777 | 0.068138 | 0.068138 | +6.32 | -6.32 | 0.000 | 1.000 |
| **Ins** | -9.736925 | 9.736925 | 1.232051 | 1.232051 | -5.41 | +5.41 | 1.000 | 0.000 |
| **NAA** | 2.575131 | -2.575131 | 0.091893 | 0.091893 | +5.32 | -5.32 | 0.000 | 1.000 |
| **Tau** | 3.049511 | -3.049511 | 0.108420 | 0.108420 | +5.55 | -5.55 | 0.000 | 1.000 |
| **PCho+GPC** | 0.030238 | -0.030238 | 0.004181 | 0.004181 | +0.46 | -0.46 | 0.323 | 0.677 |

*Supporting Table 3. Group-level statistical results for metabolite concentrations from the unpaired t-test run on the in vivo dwMRS data. WT: Wild Type, CNTF: Cytokine Ciliary Neurotrophic Factor injection.*

### Supporting Information References
